# Wnt binding to Coatomer proteins directs secretion on exosomes independently of palmitoylation

**DOI:** 10.1101/2023.05.30.542914

**Authors:** Uxia Gurriaran-Rodriguez, David Datzkiw, Leandro G. Radusky, Marie Esper, Fan Xiao, Hong Ming, Solomon Fisher, Marina A. Rojas, Yves De Repentigny, Rashmi Kothary, Adriana L. Rojas, Luis Serrano, Aitor Hierro, Michael A. Rudnicki

## Abstract

Wnt proteins are secreted hydrophobic glycoproteins that act over long distances through poorly understood mechanisms. We discovered that Wnt7a is secreted on extracellular vesicles (EVs) following muscle injury. Structural analysis identified the motif responsible for Wnt7a secretion on EVs that we term the Exosome Binding Peptide (EBP). Addition of the EBP to an unrelated protein directed secretion on EVs. Disruption of palmitoylation, knockdown of WLS, or deletion of the N-terminal signal peptide did not affect Wnt7a secretion on purified EVs. Bio-ID analysis identified Coatomer proteins as candidates responsible for loading Wnt7a onto EVs. The crystal structure of EBP bound to the COPB2 coatomer subunit, the binding thermodynamics, and mutagenesis experiments, together demonstrate that a dilysine motif in the EBP mediates binding to COPB2. Other Wnts contain functionally analogous structural motifs. Mutation of the EBP results in a significant impairment in the ability of Wnt7a to stimulate regeneration, indicating that secretion of Wnt7a on exosomes is critical for normal regeneration *in vivo*. Our studies have defined the structural mechanism that mediates binding of Wnt7a to exosomes and elucidated the singularity of long-range Wnt signalling.

## Introduction

Wnt proteins are a conserved family of secreted glycoproteins that govern essential developmental, growth, and regenerative processes, and are also involved in pathological conditions such as cancer^1^. Wnt signaling plays multiple roles in regulating stem cell function, including proliferation, cell polarity and symmetric division, motility, and fate specification^2, 3^. Despite the relative hydrophobicity due to palmitoylation^4^, Wnt proteins actively participate in long-range paracrine signaling between Wnt-producing cells and distal recipient cells^4^. Several mechanisms have been proposed to explain long-range Wnt signaling including transfer of Wnt proteins via lipoproteins^5, 6^, cell extensions called cytonemes^7^, association with soluble Wnt-binding proteins^8, 9^, or via a class of extracellular vesicle (EVs) called exosomes^10–12^.

Exosomes are 40-150 nm small EVs of endocytic origin involved in intercellular communication that transfer bioactive cargo, for example lipids, proteins, microRNAs, and mRNAs, to distal cells^13^. Several *in vitro* studies have shown different Wnt proteins are secreted on the surface of exosomes, and that exosomal-Wnts are fully capable of eliciting appropriate signaling in target cells^10–12^. Moreover, examples have been noted where significant amounts of Wnt protein are secreted on exosomes^14^. Considerable *in vivo* evidence derived from studies in *Caenorhabditis*^15^ and *Drosophila*^10, 14, 16^, support the importance of exosomes for long-range Wnt signaling, yet how Wnts are tethered to these migratory organelles remains unknown.

Following acute injury in adult skeletal muscle, Wnt7a is highly upregulated where it positively stimulates regenerative myogenesis by acting at multiple levels^17^. Wnt7a/Fzd7 signaling via the planar-cell-polarity (PCP) pathway stimulates symmetric muscle stem cell expansion^18^ and motility^19^. Wnt7a/Fzd7 signaling via the AKT/mTOR pathway in myofibers stimulates anabolic growth and hypertrophy^20^. Consequently, intramuscular injection of Wnt7a protein significantly ameliorates disease progression in *mdx* mice, a mouse model for Duchenne Muscular Dystrophy (DMD)^21^. Together, these findings indicate that Wnt7a is a promising candidate therapy for DMD. However, systemic delivery of Wnt7a via the circulation has remained a challenge because of the high hydrophobicity conferred by the conserved palmitoylation. We therefore set out to decipher the mechanism that targets Wnt7a to exosomes to provide insight into long-range Wnt signalling for the treatment of neuromuscular diseases.

Here we found that Wnt7a is secreted at high levels on exosomes following muscle injury. We performed structure function analysis and identified a novel specific signal sequence in Wnt7a that we termed the Exosome Binding Peptide (EBP). Notably, linking of the EBP sequence to other proteins resulted in their secretion on EVs. Using Bio-ID, we identified Coatomer proteins as necessary for EBP binding and the trafficking of Wnt7a to the exterior of exosomes. Finally, isothermal calorimetry (ITC) and mutagenesis confirmed that the direct interaction occurs between COPB2 and the positively charged motif KIK in the EBP. These findings elucidate the molecular mechanism for Wnt secretion on exosomes and identify a novel trafficking route that appears to function independently of the classical Wnt secretion routes.

## Results

### Muscle injury triggers secretion of Wnt7a on the surface of exosomes

Wnt7a expression is highly upregulated in newly differentiating myofibers following acute injury of skeletal muscle^22^. Examination of muscle cryosections 96 h following cardiotoxin injury by Immunogold Electron Transmission Microscopy (iTEM) labeling revealed that high levels of Wnt7a are secreted on the surface of exosomes, and stimulates muscle regeneration (Fig. 1a). Examination of Wnt7a-HA tagged (Wnt7a-Human influenza hemagglutinin) transfected HEK293T cells by iTEM revealed that Wnt7a is secreted both as free protein and on the surface of exosomes (Extended Data Fig. 1a). Importantly, Wnt7a does not colocalize with lysosomal-associated membrane protein 1 (Lamp1) or Mannose-6-phosphate receptor (M6PR), suggesting Wnt7a is not degraded in HEK293T cells (Extended Data Fig. 1b).

**Fig. 1.**
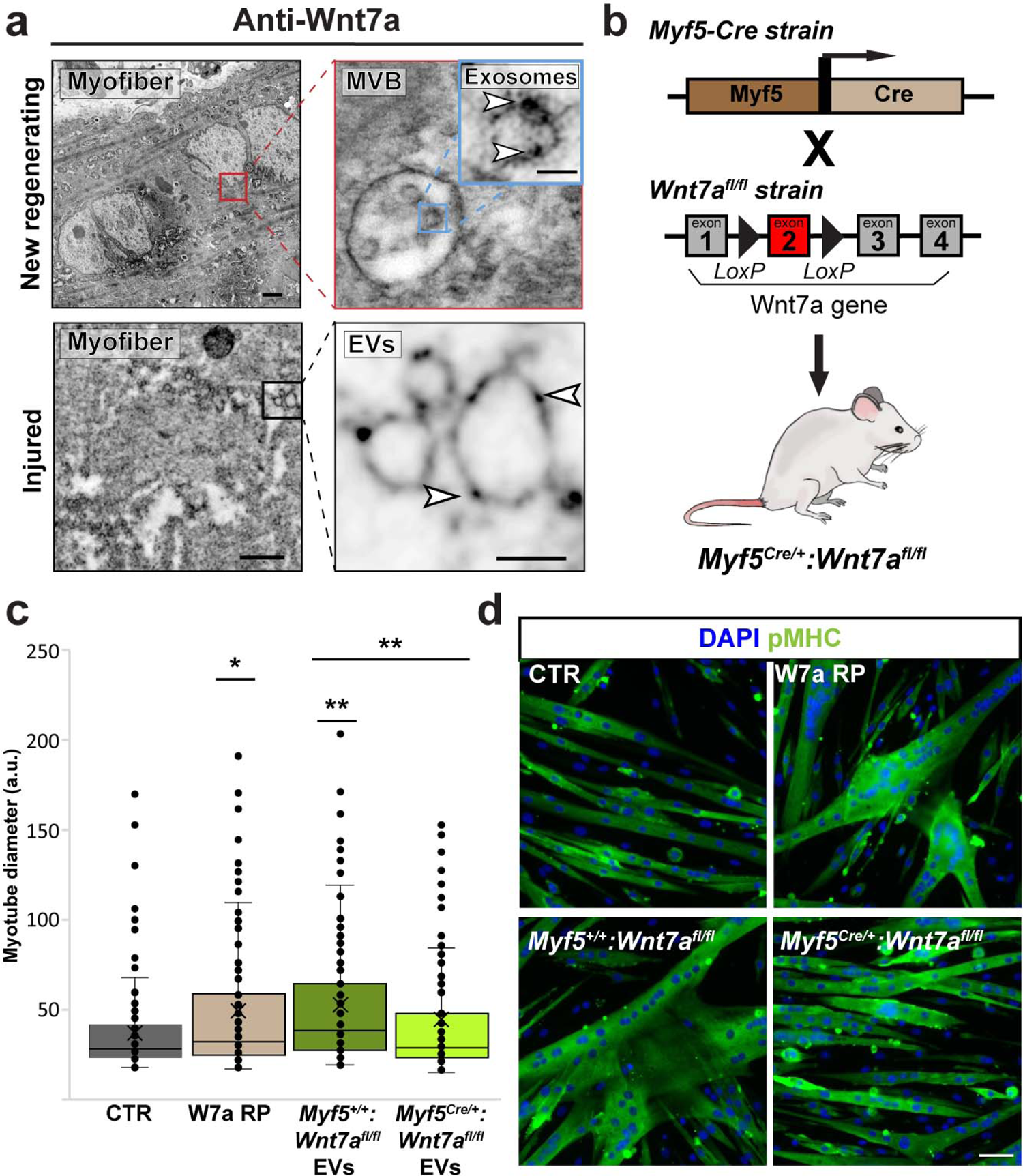
Muscle injury triggers secretion of Wnt7a on the surface of exosomes. **a**, iTEM of anti-Wnt7a labeling of new regenerating myofibers at 96 h post-CTX injury from WT mice, shows Wnt7a secretion on EVs contained in a MVB (upper panels). Scale bar 500 nm and 100 nm respectively. iTEM of anti-Wnt7a labeling of injured TA muscle from WT mice, shows the presence of Wnt7a on EVs secreted by regenerating myofibers (lower panels). Scale bar 500 nm and 100 nm respectively. **b**, Schematic representation of mouse strains used to generate conditional Wnt7a floxed in Myf5 expressing cells. **c**, Hypertrophy assay of murine primary myotubes treated with EVs from muscle decreases hypertrophy after Wnt7a deletion. Data shown as fold change of myotube diameter over the control (%); Wnt7a recombinant protein was used as a positive control. **d**, pMHC IF representative images of hypertrophied myotubes after muscle EVs stimulation containing Wnt7a (n=3). Scale bar 50 μm. Experiments are representative of n = 4 biological replicates. Experiments are representative of three independent biological replicates with 8 technical replicates each time. Immunogold Transmission electron microscopy (iTEM), Multivesicular Bodies (MVB), Extracellular Vesicles (EVs), Pan myosin heavy chain (pMHC), Tangential Flow Filtration (TFF). Statistical analysis was performed using two-sided Student’s t-test. Data are mean ± s.e.m. Source (*p<0.05, **p<0.005, ***p<0.0005).

To eliminate contamination with free secreted proteins and cytoplasmic fragments, which are typically found when using ultracentrifugation to concentrate EVs^23^, we adapted a protocol using tangential flow filtration (TFF)^24^. This approach allows simultaneous purification of both Wnt7a bound to EVs (retentate fraction) and freely secreted Wnt7a (permeate fraction) (Extended Data Fig. 1c). Quantification following TFF purification indicates that over 60% of secreted Wnt7a from HEK293T cells is bound to EVs (Extended Data Fig. 1d). Dynamic Light Scattering measurement and iTEM confirmed that the purified EVs exhibited a size distribution with the highest concentration around 150 nm, and no difference in diameter between EVs from empty vector- or Wnt7a-transfected HEK2983T cells (Extended Data Fig. 1e, f).

EVs isolated from regenerating muscle by TFF carried high levels of Wnt7a (Extended Data Fig. 2a-c). We previously established the capacity of Wnt7a to promote hypertrophy *in vitro*^25^. Notably, purified EVs from regenerating muscle readily induced hypertrophy of cultured myotubes, indicating that Wnt7a on EVs exhibits robust bioactivity (Extended Data Fig. 2d, e).

**Fig. 2.**
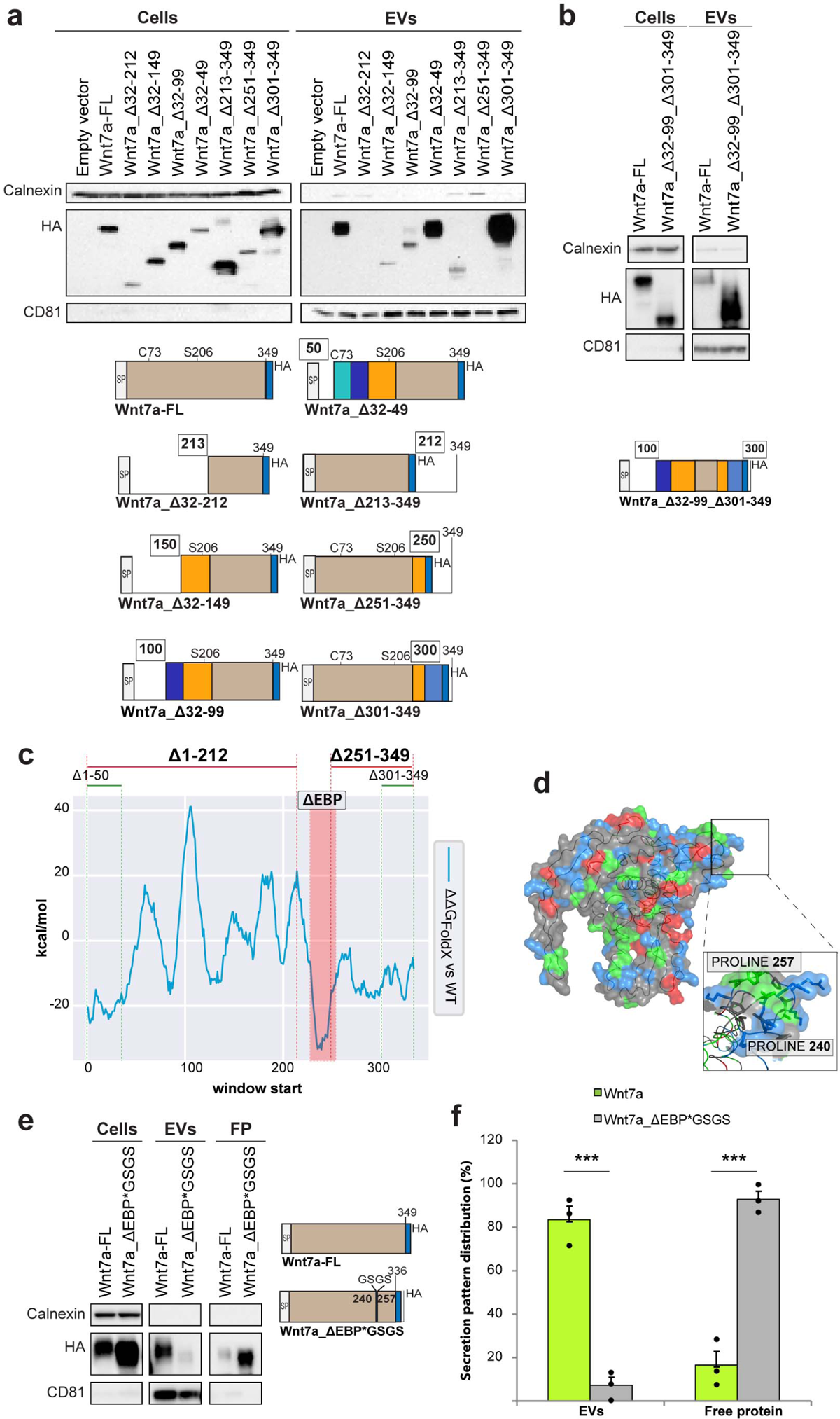
Wnt7a secretion on EVs is regulated by an internal signal peptide. **a**, Immunoblot analysis shows interruption of EV secretion following deletion beyond position 100 aa but not after position 300. **b**, Immunoblot analysis showing the internal 100-300aa sequence of Wnt7a is sufficient for EV secretion. **c**, ΔG_FoldX_ of Wnt7a indicates Δ1-49 and Δ301-349 do not affect folding ─ΔΔG < 0 respect to WT protein─ and function is not lost. Δ1-212 affects protein folding. Δ251-349 does not affect folding but function is lost since a region of the EBP is truncated. **d**, Surface of Wnt7a with negative charged residues in red, positive charged residues in blue, and hydrophobic residues in green. The EBP is positively charged. **e**, Immunoblot analysis, demonstrates replacent of EBP with a GSGS linker abrogates EV secretion in favour of free protein secretion. **f**, Quantification of Wnt7a expression on EVs and Free Protein fractions with and with-out ESP. Experiments are representative of three independent biological replicates performed in HEK293T cells transfected with different Wnt7a-HA tagged truncates. Statistical analysis was performed using two-sided Student’s t-test. Data are mean ± s.e.m. Source (***p<0.0005).

To determine whether the bioactivity of Wnt7a-EVs is due to the Wnt7a cargo, we isolated EVs from regenerating muscle from mice with a functional Wnt7a gene (*Myf5^+/+^:Wnt7a^fl/fl^*)^26^, or from mice where Wnt7a is specifically deleted in muscle (*Myf5^Cre/+^:Wnt7a^fl/fl^*), and conducted a hypertrophy assay *in vitro* (Fig. 1b). Immunofluorescence from injured *Myf5^Cre/+^:Wnt7a^fl/fl^* muscle indicates an absence of Wnt7a, whereas injured *Myf5^+/+^:Wnt7a^fl/fl^*muscle exhibits high-level expression of Wnt7a (Extended Data Fig. 2f). Accordingly, immunoblot and iTEM analysis of EVs isolated from injured *Myf5^Cre/+^:Wnt7a^fl/fl^*muscle revealed an absence of Wnt7a, whereas EVs isolated from injured *Myf5^+/+^:Wnt7a^fl/fl^*muscle exhibit the presence of high levels of Wnt7a (Extended Data Fig. 2g,h). EVs isolated from Wnt7a expressing regenerating *Myf5^+/+^:Wnt7a^fl/fl^*muscle induced a hypertrophic response in primary murine myotubes in a similar manner to recombinant Wnt7a (Fig. 1c-d). By contrast, EVs isolated from *Myf5^Cre/+^:Wnt7a^fl/fl^* regenerating muscle lacking Wnt7a did not induce significant hypertrophy (Fig. 1c-d). Therefore, the presence of Wnt7a is necessary for the hypertrophic response.

### Wnt7a secretion on exosomes is regulated by an internal binding peptide

Towards mapping the region required for secretion on exosomes, we constructed a series of N-terminal and C-terminal deletions of Wnt7a-HA (Fig. 2a). Initial N-terminal deletions were performed leaving in place the 31 aa signal peptide (SP) required for secretion of free protein^27^. The Wnt7a variants were expressed in HEK293T cells and the amount of Wnt7a secreted on EVs was assessed by immunoblot analysis (Fig. 2a and Extended Data Fig. 3a).

**Fig. 3.**
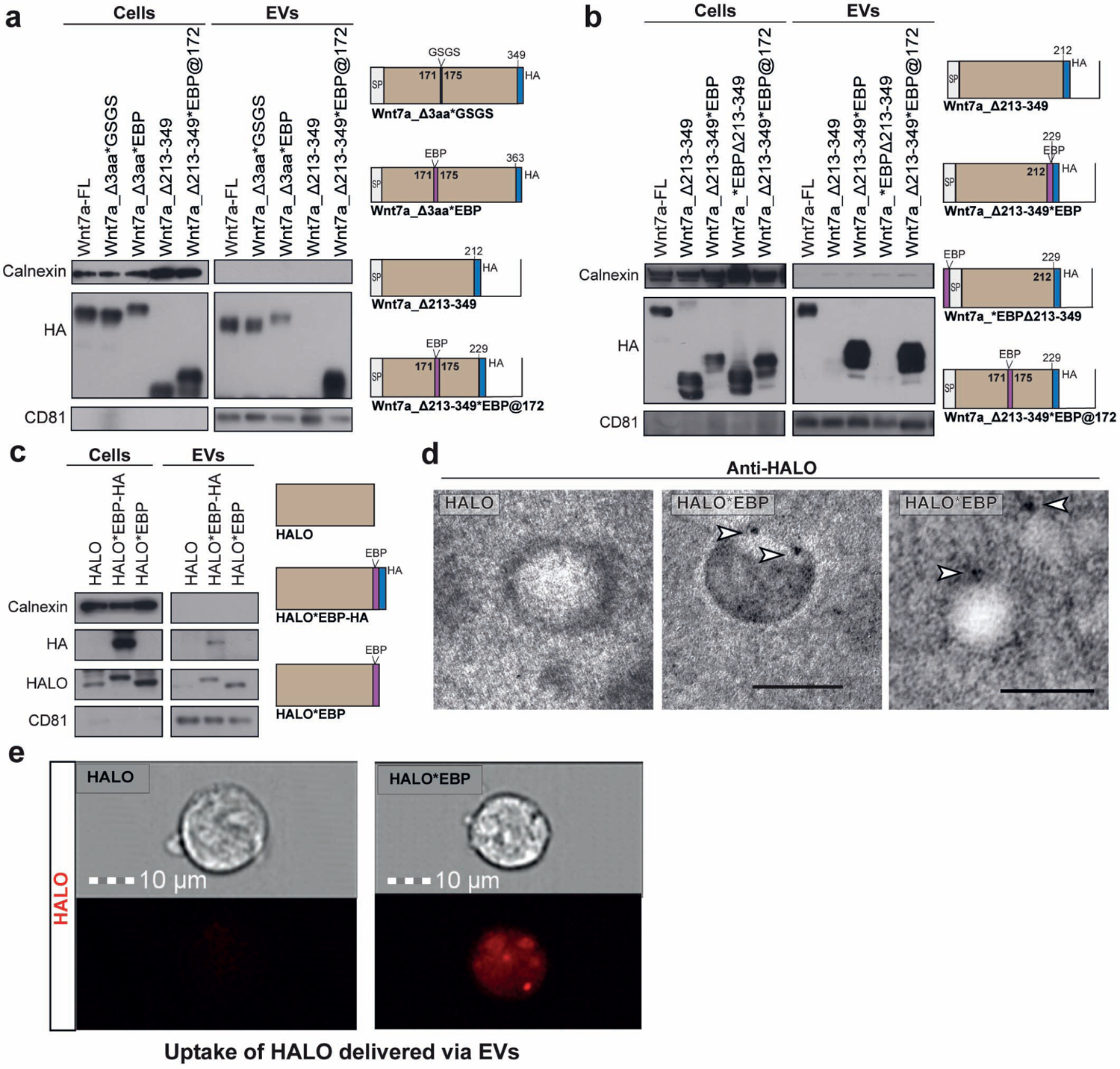
The EBP targets proteins for extracellular secretion on EVs. **a**, Restoration of EV secretion by inserting EBP into an upstream domain of Wnt7a truncate that is not secreted on EVs. The insertion does not perturb the stability of the full-length protein (Wnt7a-Δ3aa*GSG versus Wnt7a-Δ3aa*EBP). Insertion of EBP to this site (Wnt7a-Δ213-249*EBP) restores EVs localization to Wnt7a-Δ213-249. **b**, C-terminal linked EBP but not N-terminal linked, confers targeting of Wnt7a-Δ213-249 onto EVs. **c**, Linking of EBP HALO protein results in EV secretion. **d,** iTEM images of EVs with anti-HALO immunostaing showing expression of HALO*EBP on the surface of EVs. **e**, Detection of HALO by fluorescence inside HEK293T cells following treatment with HALO*EBP EVs versus HALO EVs. Experiments are representative of three independent biological replicates performed in HEK293T cells transfected with different Wnt7a-HA and HALO tagged truncates.

Wnt7a secretion on EVs was not impaired upon deletion of the N-terminal 68aa following the SP (Wnt7a_Δ32-99), and the C-terminal 48aa (Wnt7a_Δ301-349) (Fig. 2a). By contrast, deletion of additional sequences from the N-terminus (Wnt7a_Δ32-149) and C-terminus (Wnt7a_Δ251-349) appeared to abrogate secretion on EVs (Fig. 2a and Extended Data Fig. 3a). Indeed, Wnt7a lacking both the first 99aa and the last 48aa (Wnt7a_Δ1-99_Δ301-349), were readily secreted on EVs (Fig. 2b).

To identify the region that mediates the targeting to exosomes, we analyzed a 3D model of Wnt7a based on the structure of XWnt8a^28^ (Extended Data Fig. 3b). Energetic analysis after truncating successive 15 aa residue regions using FoldX (ΔG_FoldX_)^29^ revealed that the deletion of amino acids between positions 240 and 257 does not interfere with Wnt7a structural folding stability (Fig. 2c). This low energetic region is a result of a hydrophobic random coil structure flanked by two prolines between position 240 and 257 (Fig. 2d).

The region was then investigated as a potential binding site that would mediate targeting of Wnt7a to exosomes. Replacement of the 18 aa sequence between position 240 and 257 with the linker domain GSGS (Wnt7a_ΔEBP*GSGS), resulted in a loss of Wnt7a targeting to EVs with a corresponding increase in secretion of free Wnt7a protein, and with no effect on total Wnt7a protein expression (Fig. 2e-f). Therefore, the sequence PVRASRNKRPTFLKIKKP, which we term the Exosome Binding Peptide (EBP), is required for targeting Wnt7a to exosomes.

### The EBP is sufficient to confer targeting to exosomes

We next investigated whether the EBP is capable of targeting proteins for secretion on exosomes. We first added the EBP to a truncated Wnt7a that we previously found to not localize to EVs (Wnt7a_Δ213-349) (Fig. 2a). We chose a specific insertion site for the EBP within the Wnt7a_Δ213-349 truncate to avoid any conformational disruption or EBP offshoring. Energetic and conformational studies with FoldX showed a loop starting at position 172 as a potential insertion site for the EBP with a similar distance between loop terminals (10.89Å versus 8.69Å), and proximal in the 3D space to the original EBP location (Extended Data Fig. 4a).

**Fig. 4.**
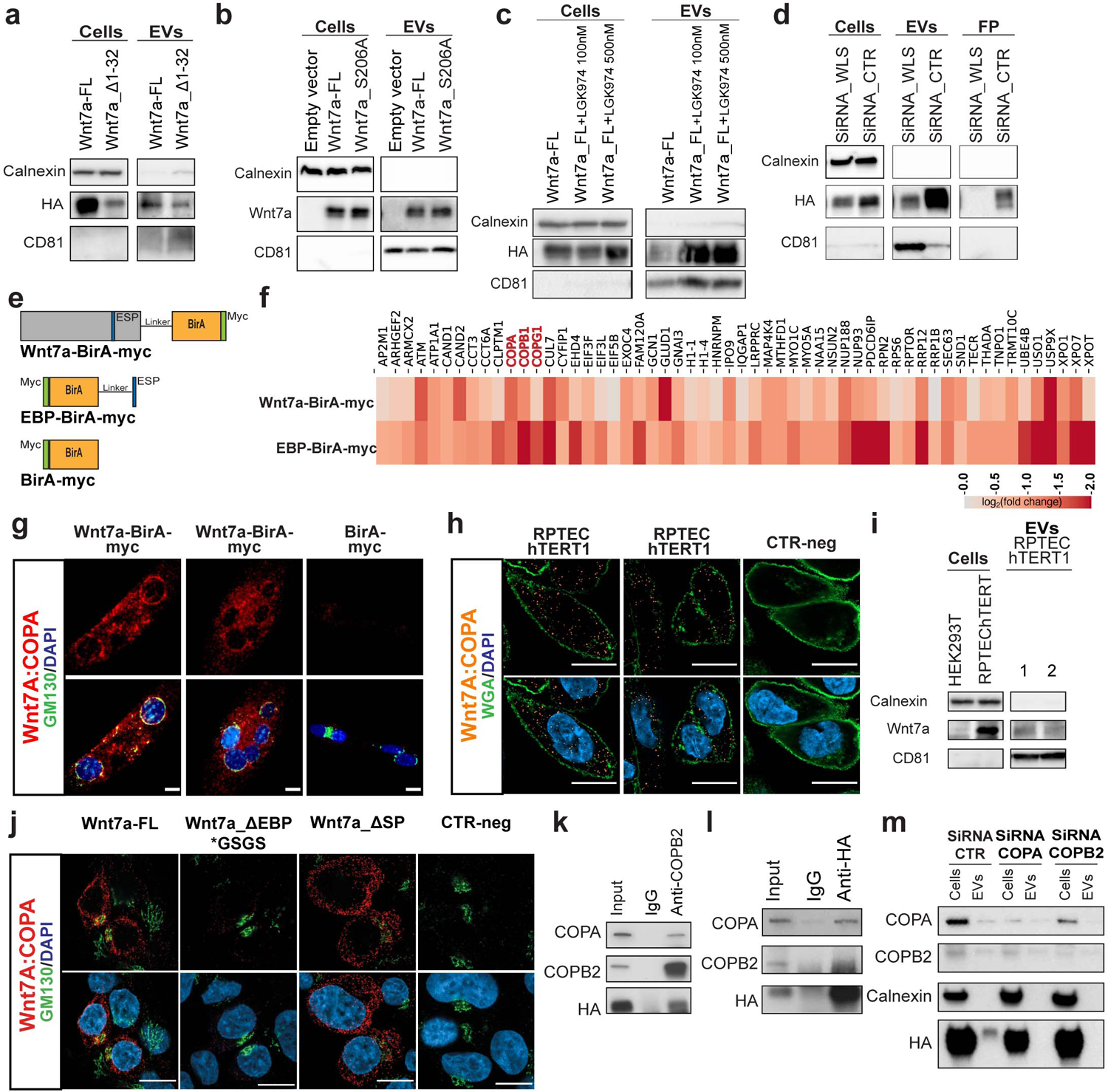
Secretion of Wnt7a on exosomes is regulated by interaction with Coatomer proteins. **a**, Signal peptide is not required for EVs-Wnt7a secretion in transfected HEK293T cells. **b,** Mutant Wnt7a_S206A, lacking the palmitoylation site, is secreted on EVs in transfected HEK293T cells. **c,** Drug inhibition of PORCN does not affect secretion of Wnt7a on EVs in transfected HEK293T cells. **d,** Knockdown of WLS in siRNA in transfected HEK293T cells does not affect secretin of Wnt7a on EVs but does abolish secretion of free Wnt7a. **e,** BirA constructs designed for the BioID analysis. **f**, Heat map displaying fold change (log2 scale) of enriched proteins in mass spectrometry versus control conditions (ESP_BirA:BirA and Wnt7a_BirA:BirA). Shown are proteins that present a minimum enrichment of 50% (log2 (FC)>0.5849) on EBP and a positive enrichment (log2 (FC)>0) on Wnt7a. COPI complex subunits are highlighted in red. **g**, Wnt7a: COPA PLA (red) performed in murine primary myotubes either expressing Wnt7a-BirA or BirA. PLA signal was counterstained with GM310 (green), a Golgi Apparatus marker, and with DAPI (blue). Scale bar 10 μm. **h**, Wnt7a:COPA PLA (orange) performed in RPTEC-hTERT1 cells. PLA signal was counterstained with WGA (green), a cell membrane marker and with DAPI (blue), showing interaction in the cytosol. CTR-neg is an internal control without Wnt7a antibody. Scale bar 10 μm. **i,** Wnt7a is secreted on EVs derived from RPTEC-hTERT1 cells. HEK293T cells were used as a negative control for Wnt7a expression. **j,** Wnt7a:COPA PLA (red) performed in HEK293T cells either expressing Wnt7a-FL, Wnt7a_ΔEBP*GSGS or Wnt7a_ΔSP. PLA signal was counterstained with GM310 (green), a Golgi Apparatus marker, and with DAPI (blue). Scale bar 10 μm. CTR-neg is an internal control without Wnt7a antibody. **k**, HEK293T cells overexpressing Wnt7a-HA were immunoprecipitated with COPB2 antibody. Wnt7a-HA interacts with COPA and COPB2. **l**, HEK293T cells overexpressing Wnt7a-HA were immunoprecipitated with HA antibody. Wnt7a-HA interacts with COPA and COPB2. **m**, Immunoblot EVs secretion analysis of Wnt7a after siRNA knock down of COPA and COPB2 shows disruption of Wnt7a-EVs secretion. Experiments are representative of three independent biological replicates.

Insertion of the linker GSG or EBP between position 171 and 175 into full length Wnt7a-HA (Wnt7a-FL) had no effect on secretion on EVs (Fig. 3a). Notably, insertion of the EBP between position 171 and 175 into Wnt7a_Δ213-349 (Wnt7a_Δ213-349*EBP@172) fully restored secretion on EVs (Fig. 3a). Moreover, addition of EBP to the C-terminus of Wnt7a_Δ213-349 (Wnt7a_Δ213-349*EBP@212) confirmed the structurally independent capacity of EBP to target proteins for secretion on EVs (Fig. 3b). By contrast, addition of the EBP to the N-terminus of Wnt7a_Δ213-349 adjacent the SP (Wnt7a_*EBPΔ213-349), did not result in secretion on EVs (Fig. 3b) suggesting that proximity of both signal peptides interferes with EV targeting.

We next asked whether linking the EBP to a non-Wnt protein would direct secretion on exosomes. Therefore, the EBP was fused to the HALO tag, a 297 aa peptide derived from a bacterial enzyme designed to covalently bind fluorescent ligands^30^ (Fig. 3c). The HALO protein alone was not secreted on EVs, whereas HALO*EBP-HA and HALO*EBP were both efficiently secreted on EVs (Fig. 3c). Detection of HALO using iTEM confirmed the presence of HALO*EBP on the surface of EVs (Fig. 3d). Furthermore, purified EVs efficiently delivered the EBP-tagged HALO protein to recipient HEK293T cells, as assessed by labeling EVs with a specific fluorescent tag for HALO (Fig. 3e and Extended Data Fig. 4b-c). By contrast, EVs isolated from HALO overexpressing cells did not deliver HALO to recipient cells as revealed by the absence of fluorescence staining (Fig. 3f). Therefore, we conclude that the EBP is sufficient to mediate targeting of proteins to exosomes that can then be delivered to recipient cells.

### Wnt7a secretion on exosomes is not dependent on the signal peptide, palmitoylation, or WLS binding

To investigate the involvement of the ER-Golgi secretory pathway, we assessed the requirement of several mechanisms that have been attributed to Wnt protein secretion. Interestingly, we found that deletion of the N-terminal SP did not alter the ability of Wnt7a to be secreted on EVs (Fig. 4a). Furthermore, deletion of the SP from the minimal Wnt7a_Δ1-99_Δ301-349 (Figure 2b) construct did not affect secretion on EVs (Extended Data Fig. 5a). Notably, Wnt7a_Δ1-99_Δ301-349 showed over 100% increase of secretion on EVs than its counterpart Wnt7a_Δ32-99_Δ301-349 with the SP (Extended Data Fig. 5b). Lastly, all versions of Wnt7a that were previously tested (Fig. 2a), are fully secreted on EVs in absence of SP (Extended Data Fig. 5c).

**Fig. 5.**
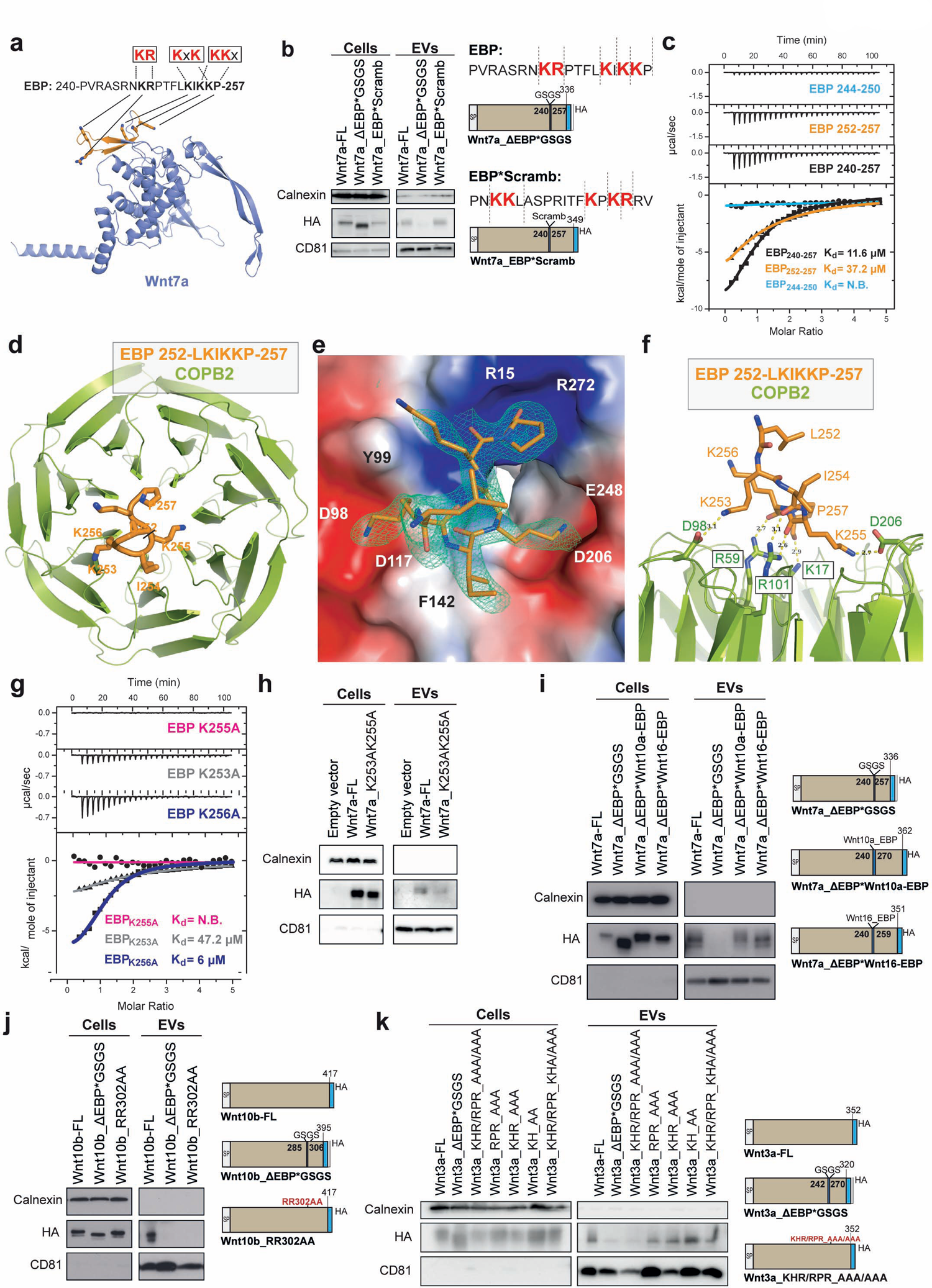
The KIK motif mediates binding to Coatomer proteins. **a**, Predicted Wnt7a structure in AlphaFold (blue) with the EBP region highlighted in orange and the three positively charged motifs (red) within the EBP. Note that EBP is a solvent-exposed region. **b,** Wnt7a-ESP*Scramble mutant maintains the dilysine motif and exhibits no impairment in EV secretion. **c,** Isothermal calorimetry measurements for COPB2_1-304_ binding to potential dilysine/arginine motifs within the EBP. Wild-type COPB2_1-304_ binds to the LKIKKP sub-region. **d-f,** views of the KxKx motif of Wnt7a bound to COPB2_1-304_. **d**, Top view of the WD-repeat domain of COPB_1-304_ (green) with the LKIKKP peptide (orange) in ribbon representation. **e,** Close-up view of the LKIKKP peptide with a difference electron density map calculated by omitting the peptide and contoured at 3σ (blue mesh). COPB2 surface is colored by electrostatic potential ranging from −5kT/e (red) to 5kT/e (blue). **f,** Lateral view of the binding motif with hydrogen bonds and distances. **g,** Structure-based point mutations confirm the molecular recognition of the KxKx motif in ITC assays. **h**, Double lysine mutation of K253 and K255 by alanine disrupts Wnt7a-EVs secretion. **i**, Replacement of Wnt7a-EBP by either Wnt10a-EBP or Wnt16 EBP containing KR and RR (right panel). Replacement with Wnt10a-EBP or Wnt16 EBP rescues Wnt7a-EVs secretion. **j**, EV secretion analysis of Wnt10b after EBP removal or double arginine mutation within its EBP (right panel). Double arginine mutation disrupts secretion of Wnt10b on EVs to the same extent as removal of the entire Wnt10b EBP sequence. **k**, Secretion analysis of Wnt3a after EBP removal or mutation of the entire positively charged motifs RPR, KHR, KH within its EBP (right panel). Only concomitant mutation of RPR and KHR motifs disrupts secretion of Wnt3a on EVS to the same extent as removal of the entire Wnt3a EBP sequence. Experiments are representative of three independent biological replicates performed in HEK293T cells transfected with different Wnt-HA tagged truncates.

Several groups have asserted the importance of palmitoylation of Wnt for secretion and activity^31–33^. Therefore, we tested whether Wnt7a was secreted on EVs following mutation of the palmitoylation site at serine 206^34^. Previously, we noted that palmitoylation is not required for Wnt7a secretion or activity^35^. By contrast, palmitoylation is critical for Wnt3a secretion^2, 36^. Notably, we observed that secretion of Wnt7a on EVs was unaffected by mutation of the palmitoylation site (Fig. 4b). Similarly, inhibition with LGK974, a chemical inhibitor of PORCN^37^, had no apparent effect on Wnt7a secretion on EVs (Fig. 4c). Therefore, we conclude that palmitoylation does not play a role in regulating the secretion of Wnt7a on exosomes.

WLS is a chaperone transmembrane protein that binds Wnt to direct secretion at the cell membrane^38, 39^. Our data confirmed that secretion of Wnt7a free protein was entirely abolished upon siRNA knockdown of WLS (Extended Data Fig. 5d). By contrast, Wnt7a secretion on EVs was readily detectable following siRNA knockdown of WLS (Fig. 4c). Therefore, we conclude that WLS is not required for Wnt7a-secretion on exosomes.

### Wnt7a binds Coatomer proteins

To elucidate the molecular basis whereby the EBP mediates the secretion of Wnt7a on exosomes, we performed BioID interactome analysis to identify candidate binding proteins. We linked BirA, a highly efficient proximity dependent biotin ligase that biotinylates proteins in close proximity to the bait proteins (Extended Data Fig. 5e)^40^. Mouse primary myoblasts were generated to express myc-tagged Wnt7a-BirA, ESP-BirA, or BirA (Fig. 4e and Extended Data Fig. 5f). In addition, the ability of Wnt7a-BirA-myc to be localized to the myoblasts-derived EVs was verified by mass spectrometry (Extended Data Fig. 5g). Moreover, iTEM revealed localization of Wnt7a and the Myc tag on the EV surface regardless of permeabilization (Extended Data Fig 5h, i).

Mass spectrometric identification of biotinylated proteins isolated from transfected primary myoblasts revealed that Coatomer proteins COPA and COPB2 were among the most enriched candidates within the EBP-BirA and Wnt7a-BirA interactomes (Fig. 4f). Indeed, GO-term analysis of the interactome indicated that Coatomer proteins were the amongst the highest represented hits (Extended Data Fig. 6a). Importantly, COPA and COPB2 have been previously noted to be present at high levels on EVs^41, 42^. Proximity Ligation Assays (PLA) between Wnt7a and COPA or COPB2 in myotubes from murine primary myoblast, confirmed the interaction of Wnt7a with these Coatomer components (Fig. 4g and Extended Data Fig. 6b respectively). Interestingly, PLA detected the interaction between Wnt7a and COPA primarily at the Golgi whereas the interaction between Wnt7a and COPB2 appeared enriched proximal to the cell membrane (Fig. 4f and Extended Data Fig. 6b respectively), consistent with the suggestion that different coatomer proteins have distinct roles in endosomes and MVBs as previously described^43, 44^.

**Fig. 6.**
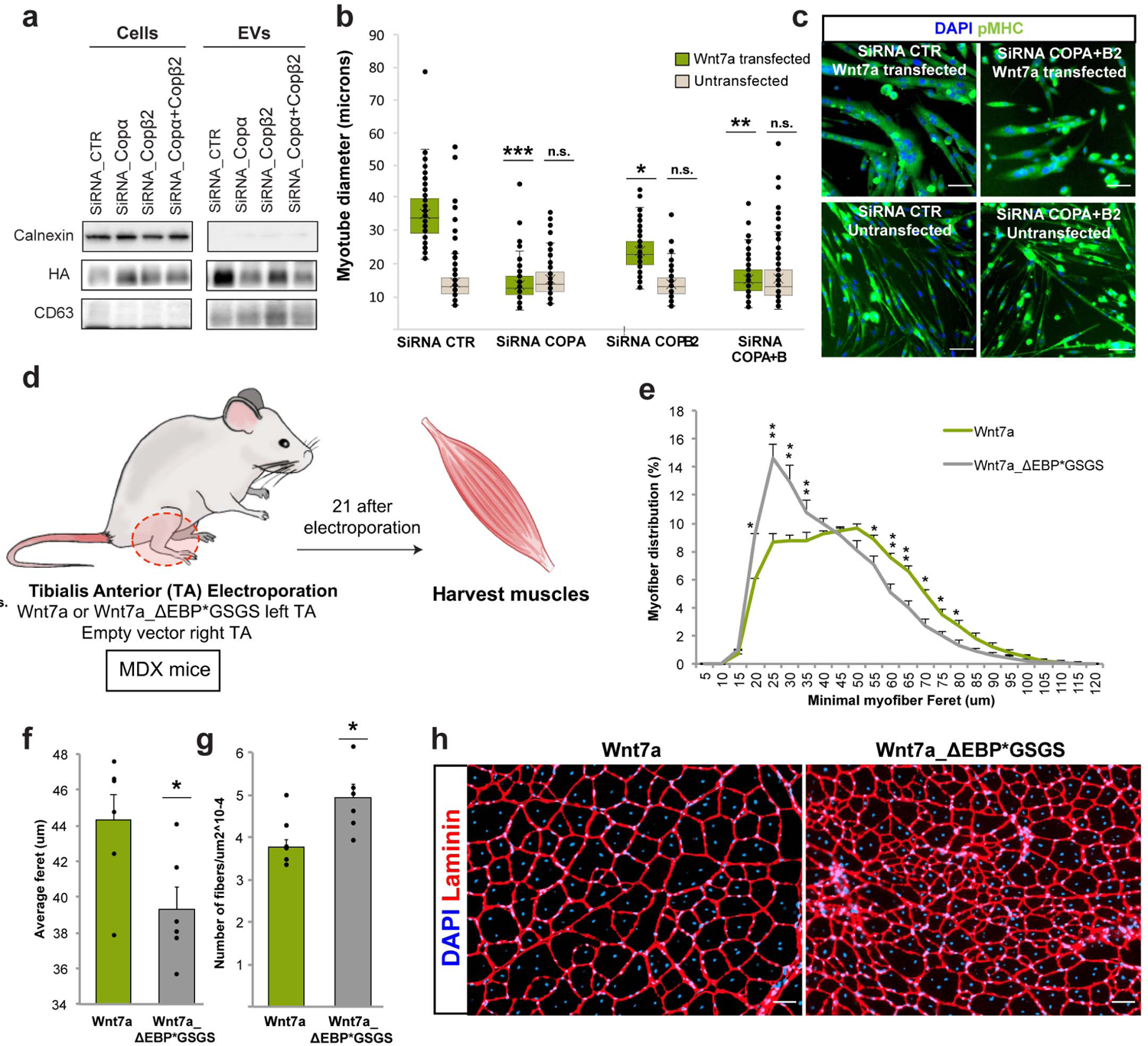
EBP sequence of Wnt7a is required for stimulation of muscle regeneration. **a**, Knockdown of COPA, COPB2 using siRNA disrupts Wnt7a-EVs secretion. **b**, Knock down of COPA, COPB2 decreases hypertrophy in Wnt7a transfected myotubes. Conversely, knock down did not affect hypertrophy in myotubes not expressing Wnt7a. Data shown as fold change of myotube diameter over the siRNA control (%). **c**, Representative images after simultaneously siRNA knock down of COPA and COPB2 in myotubes. Scale bar 50 μm. **d**, Schematic representation of *in vivo* workflow. **d**, Myofiber caliber distribution comparing TA electroporated with Wnt7a (green) vs Wnt7a_ΔEBP*GSGS (grey). **f,** Average feret comparing TA electroporated with Wnt7a (green) vs Wnt7a_ΔEBP*GSGS (grey). **g,** Quantification of fiber number comparing TA electroporated with Wnt7a (green) vs Wnt7a_ΔEBP*GSGS (grey). **h,** Section of TA muscles showing reduced myofiber caliber after electroporation of Wnt7a_ΔEBP*GSGS into TA muscle of *mdx* mice. Scale bar 100 μm. Pan myosin heavy chain (pMHC). *Tibialis anterior* (TA). *In vitro* experiments are representative of three independent biological replicates performed in murine primary myoblasts. *In vivo* experiments are representative of six mice per condition. Statistical analysis was performed using two-sided Student’s t-test. Data are mean ± s.e.m. Source (*p<0.05, **p<0.005).

These findings were corroborated in a human epithelial immortalized RPTEC-hTERT1 cells derived from kidney cortex that endogenously expresses Wnt7a (Extended Data Fig. 6c-d). PLA between Wnt7a and COPA revealed the interaction of endogenous Wnt7a with COPA in the cytoplasm (Fig.4h). Immunoblot data demonstrate that RPTEC-hTERT1 cells endogenously express Wnt7a that is secreted on EVs (Fig. 4i). Therefore, interaction of Wnt7a with Coatomer and secretion on exosomes is not a consequence of overexpression.

These results were also verified in HEK293T cells. PLA did not detect an interaction between Wnt7a_ΔEBP*GSGS (lacking the EBP), and COPA, whereas the interaction was detected between Wnt7a_ΔSP (lacking the N-terminal signal peptide), and COPA (Fig. 4j and Extended Data Fig. 6e). Reciprocal immunoprecipitation-western blot analysis suggested that Wnt7a interacts with COPA and COPB2 in the same complex (Fig. 4k, l). Mass Spectrometry analysis of lysates of Wnt7a-HA expressing EVs immunoprecipitated with anti-HA antibody confirmed the interaction of COPA and COPB2 with Wnt7a in EVs (Extended Data Fig. 6f). To assess the requirement of COPA and COPB2 for Wnt7a secretion on exosomes, siRNA knockdown of COPA and COPB2 was performed (Fig. 4m). Notably, knock down of COPA or COPB2 resulted in an abrogation of Wnt7a secretion on EVs (Fig. 4m). Together, these results suggest a novel secretion mechanism whereby Wnt7a trafficking to exosomes does not require the N-terminal signal peptide (SP) or palmitoylation and occurs through an interaction with the Coatomer proteins.

### The KxK motif mediates EBP binding to the Coatomer

Polypeptides containing the positively charged motifs KKxx, KxKxx, or RKxx have been previously shown to bind COPA and COPB2^45^. Therefore, we evaluated whether the KR, KIK and KKP motifs within the EBP are required for secretion of Wnt7a on exosomes (Fig. 5a). Replacing the EBP with a scrambled sequence (PN**KKL**ASPRITF**KPKR**RV), which maintains the positively charged motifs (Wnt7a_EBP*Scramb), had no effect on Wnt7a-EVs secretion in HEK293T cells (Fig. 5b).

Isothermal titration calorimetry (ITC) confirmed high affinity binding of the 18 aa EBP polypeptide with the N-terminal WD-repeat of COPB2 with a dissociation constant (Kd) of 11.6 μM (Fig. 5c). The C-terminal half of the polypeptide (252-257) bound to COPB2 with a Kd of 37.2 μM. By contrast, the N-terminal half (244-250) did not bind COPB2. These data support the notion that the KIKK motif mediates binding to COPB2 (Fig. 5c, and Extended Table 1).

To elucidate the molecular details for the interaction, we solved the structure of the EBP (252-257) bound to the COPB2 at 1.8 Å resolution by X-ray crystallography (Fig. 5d-f and Extended Data Table 2). The interaction involves two main contacts of the lysine side chains K253 and K255 with two acidic patches at the central opening of the β-propeller top face (Fig. 5e). More specifically, the amino groups of K253 and K255 interact with the carboxylate groups of D98 and D206 within the WD-repeat domain (S. cerevisiae COPB2 numbering) (Fig. 5f). In addition to the two primary contacts, a secondary contact involves the backbone carbonyl oxygen atoms K255, K256 and P257 of EBP, which interact with the guanidinium groups of R59, R101 and the amino group of K17 in the WD-repeat domain (Fig. 5f). The binding mode of EBP to COPB2 is very similar to that of the KxKxx motif of yeast Emp47p, with sequence IKTKLL (Extended Data Fig. 7a), and with comparable affinity (Kd = 49 μM for Emp47p)^46^.

We tested our model in solution using a combination of ITC and structure-guided mutants. As expected, K255A or K253A substitutions drastically reduced or even abolished the ESP-COPB2 interaction, whereas K256A substitution, which is not involved in COPB2 binding, did not significantly affect the affinity (Fig. 5g). To confirm the ITC results we transfected HEK293T cells with a double point mutation of the lysines residues to alanine K253 and K255 (Fig. 5h). Immunoblot data corroborated that disruption of the KIK motif blocks the secretion of Wnt7a on EVs. These data together confirm that COPB2 and COPA regulate Wnt7a trafficking to exosomes by interacting with the KIK motif within the EBP (Fig. 5h).

The EBP appears to function as a linking peptide that connects the N- and C-terminal domains of the 19 human Wnt proteins with a sequence that is highly variable in length and aa composition (Extended Data Fig. 8a). Notably, the KK motif is present in Wnt2 (Extended Data Fig. 8a), and other positively charged motifs, such as KR, are present in Wnt5b, Wnt8a, Wnt11 and Wnt16. Moreover, another positively charged motif, RR, is present in Wnt2b, Wnt4, Wnt10a, Wnt10b and Wnt16 proteins (Extended Data Fig. 8a). Together, this suggests the possibility of a conserved mechanism mediating secretion on exosomes across the entire Wnt family (Extended Data Fig. 8a)

To test the ability of candidate EBPs from different Wnts to mediate secretion on exosomes, we transfected HEK293T cells with a mutant where the EBP of Wnt7a was replaced with either the EBP from Wnt10a, containing only the RR motif, or the EBP from Wnt16, that contains both motifs RR and KR (Extended Data Fig. 8a). Both the Wnt10a and the Wnt16 EBP were conferred efficient secretion on EVs (Fig. 5i). Furthermore, deletion of the EBP from Wnt10b-EVs, or double mutation of its RR motif, completely abrogated secretion of Wnt10b on EVs (Fig. 5j). Similar results were obtained in HEK293T cells with Wnt3a that contains three positively charged motifs RPR, KHR and KH. Point mutations of these three motifs separately showed no abrogation of Wnt3a secretion (Fig. 5k). Conversely, mutation of the three motifs together resulted in the loss of Wnt3a secretion on EVs, as does deletion of the entire EBP in Wnt3a (Fig. 5k).

The Wnt exosomal secretory mechanism appears to be evolutionarily well conserved. Wg the Wnt ortholog in *Drosophila Melanogaster*, contains a similar disordered loop in the linker region that contains KR and KK motifs (Extended Data Fig. 8b). Moreover, Drosophila Wg has been found to be secreted on exosomes^10, 14, 16^. Together these results suggest that the direct binding of Coatomer with Wnt family members via positively charged motifs, within the EBP domain, represents a conserved mechanism that mediates the secretion and localization of Wnts on the surface of exosomes.

### Wnt7a interaction with Coatomer is required for the muscle regenerative response

To investigate the role of Wnt7a-Coatomer interaction during the muscle regeneration we first tested Wnt7a secretion on exosomes in cultured primary myotubes following siRNA-mediated knock down of COPA or COPB2 (Extended Data Fig. 9a). We observed strongly reduced Wnt7a secretion on EVs following knock down of COPA or COPB2 (Fig. 6a). Correspondingly, COPA and COPB2 knockdown in Wnt7a expressing primary myotubes abrogated the hypertrophy response induced by Wnt7a (Fig. 6 b,c). No reduction in myotube diameter was observed following COPA or COPB2 knockdown in untransfected myotubes that do not normally express Wnt7a (Fig. 6b, c). No additive result was found when both COPA and COPB2 were knocked down, suggesting that both proteins are indispensable for Wnt7a secretion on EVs (Fig. 6b, c).

To assess the requirement for the binding of Wnt7a with coatomer for muscle regeneration *in vivo,* we electroporated the *Tibialis Anterior* (TA) muscle of male *mdx* mice with vectors expressing either full-length Wnt7a (Wnt7a-FL) or Wnt7a with the EBP replaced with a GSGS linker (Wnt7a_ΔEBP*GSGS) (Fig. 6 d), and analyzed after 21 days. Previously, we found that expression of Wnt7a in the TA muscle of *mdx* mice significantly increased myofiber caliber^47^. Electroporation of the Wnt7a-FL expression vector induced the formation of larger caliber myofibers, whereas electroporation of vectors expressing Wnt7a_ΔEBP*GSGS led to an increase in numbers of myofibers that were smaller in caliber (Fig. 6 e, f, g, h and Extended Data Fig. 9b,c). Together, these data indicate that Wnt7a secretion on EVs is required for Wnt7a bioactivity in regenerative myogenesis.

## Discussion

We have discovered a new role for coatomer proteins as mediators of Wnt7a secretion on exosomes. We discovered that a dilysine motif in a 18 aa linker domain, termed the EBP, within Wnt7a mediates binding to COPA and COPB2, and have described the structural basis of this interaction. We further show that EBP binding to COPA and COPB2 is necessary and sufficient for secretion of proteins on EVs. Notably, the EBP appears to have a conserved function across the Wnt family. Our experiments indicate that Wnt7a secretion on exosomes is required for Wnt7a bioactivity in regenerative myogenesis. Furthermore, our data is consistent with the conclusion that Wnts are displayed on the surface of exosomes via the binding to the Coatomer proteins.

Coatomer proteins have being previously involved on endosomal trafficking and biogenesis of Multivesicular Bodies (MVBs)^50^, as well as being highly enriched in EVs^51, 52^. Notably, knockdown of the orthologues of COPA or COPB2 (Dmel\αCOP and Dmel\βCOP respectively) in *Drosophila* results in adult flies that display notched wings, consistent with the notion that coatomer proteins have an essential role in Wg secretion^53^. Interestingly, Wnt5a directs the assembly of the Wnt-receptor-actin-myosin-polarity (WRAMP) structure involved in motility, and associates with MVBs and coatomer proteins^54, 55^. Moreover, downregulation of COPB2 by MicroRNA-4461 inhibits tumorigenesis derived by exosomes in colorectal cancer^56^. Therefore, Coatomer proteins may be implicated in additional cell functions and pathologies other than coating COPI vesicles for retrograde transport.

The 18 aa EBP region has been implicated in other Wnt7a functions. For example, the Reck receptor binds Wnt7a through sequences in the EBP to form a signalosome that induces canonical Wnt7a signaling^57, 58^. In addition, Wnt7a binds the canonical Frizzled co-receptor LRP6 through the EBP sequence^59^. Notably, we did not detect Reck in our BioID assays and none of the six critical aminoacids within the linker region that bind Reck are involved in the Wnt7a-Coatomer interaction. Together these findings suggest that this unstructured loop acts as an intrinsically disordered protein sequence^60^ to coordinate distinct cell-type specific functions.

Several groups have shown that Wnt secretion requires an interaction with WLS, a chaperone transmembrane protein that facilitates the secretion to the membrane^38, 39^. Moreover, WLS interacts with coatomer within COPI vesicles to mediate the recycling of WLS and thus promoting Wnt secretion^61^. This data can be interpreted to suggest that WLS acts as the linker between Coatomer and Wnt facilitating the transfer to the membrane. However, it was recently shown that the interaction between Wnt and WLS is through three hairpins where Wnts are palmitoylated and does not involve the EBP^62^. Accordingly, we found that knock down of WLS did not affect Wnt7a secretion on EVs but does result in loss of Wnt7a free protein secretion. Moreover, we did not detect any interaction of Wnt7a with WLS by Bio-ID. Consistent with our results, WntD has been observed to be active and secreted without palmitoylation^63^, and mutation of the Wnt3a palmitoylation sites only partially abrogates secretion^64^. Accordingly, we found that neither mutation of the Wnt7a palmitoylation site nor inhibition of PORCN had any effect on Wnt7a secretion on EVs.

Our experiments indicate that the N-terminal SP is not required for Wnt7a exosomal secretion, suggesting that Wnt7a trafficking onto exosomes occurs inside of the cell and not by endocytosis of free Wnt7a protein. Notably, we found that deletion of the SP enhances secretion of Wnt7a on EVs compared to its counterpart with a SP sequence. Notably, mutation of the EBP markedly enhances secretion of free protein and abrogation of secretion of Wnt on exosomes whereas inhibiting palmitoylation did the opposite. Thus, our discovery suggests that the different modes of Wnt secretion can be be modulated. Controlling the modality of Wnt secretion will enable more precise therapies for Wnt targeting as we have seen for Wnt treatment in neuronal disorders^65^.

In conclusion, we have discovered a novel role for Coatomer components in the secretion of Wnts on exosomes. Our results suggest the existence of a Wnt7a exosomal secretion pathway that functions in parallel to the ER-Golgi classical pathway, and is independent of palmitoylation. We have defined the sequence requirements for Wnt-Coatomer interaction and shown that a similar mechanism is involved in EVs secretion of multiple Wnts. Moreover, our experiments suggest that the physiological secretion of Wnt7a *in vivo* is primarily on exosomes elucidating the ability of Wnts to signal over long distances *in vivo*. Our experiments suggest that systemic delivery of Wnt7a loaded on exosomes represents a potential therapy for treating neuromuscular diseases. Importantly, the use of the EBP to direct the presentation of cargo proteins on the surface of exosomes opens the door for multiple therapeutic applications. We anticipate that our discovery will be a starting point for more sophisticated delivery systems as well as providing important insight towards understanding Wnt secretion in pathological contexts.

## Methods

### Cell culture

HEK293T cells were obtained from ATCC (CRL-3216) and verified to be free from mycoplasma contamination using the MycoSensor PCR Assay Kit (Agilent Technologies). Cells were cultured as in DMEM (Lonza) supplemented with 10% FBS, 100 U/mL penicillin, 100 U/mL streptomycin and maintained at 37°C in a humidified incubator equilibrated with 5% CO2. Wild type primary myoblasts were purified from C57BL/10ScSn M1 male mice by magnetic cell separation (MACS) as previously described by Sincennes et al^66^. Primary myoblasts were cultured on collagen-coated dishes with HAM F12-X, 10% FBS, 100 U/mL penicillin, 100 U/mL streptomycin and maintained at 37°C in a humidified incubator equilibrated with 5% CO2. For differentiation, myoblasts were grown up to 80% confluence and growth media was replaced with differentiation medium [HAM F12-X: DMEM (1:1), 5% HS, 100 U/mL penicillin, and 100 U/mL streptomycin] for 4d unless otherwise stated. During differentiation serums were treated to be free of extracellular vesicles prior to assays^67^. Wnt7a expressing primary myoblasts were obtained upon infection of primary myoblasts from C57BL/10ScSn M1 male mice, previously isolated as Sincennes et al^66^ described. Briefly, myoblasts passage 4 were seeded in 6-well plate (1.0 × 105 cells/well/4ml media) and infected with lenti-III-Ubc-Wnt7a-HA lentivirus (30ul virus/well) created in our laboratory, empty vector lentivirus as control, containing 6ug/ml Polybrene in 1.5ml culture media (Collagen-coated dishes with HAM F12-X, 10% FBS, 100 U/mL penicillin, 100 U/mL streptomycin and maintained at 37°C in a humidified incubator equilibrated with 5% CO2). Selection was done with Puromycin (2.5 ug/mL) to establish the stable cell line. Six different colonies were picked and Wnt7a expression was verified by immunoblot.

RPTEC hTERT1 cells were culture in DMEM: F12 Medium from ATCC plus the hTERT RPTEC Growth Kit from ATCC, G418 for immortalization selection, 100 U/mL penicillin, 100 U/mL streptomycin and maintained at 37°C in a humidified incubator equilibrated with 5% CO2.

### Mice and animal care

All experimental protocols for mice used in this study were approved by the University of Ottawa Animal Care Committee, which is based on the guidelines of the Canadian Council on Animal Care. Food and water were administered *ad libitum*. Muscle regeneration experiments in Fig. 1a were assessed in 8-week-old male C57BL/10ScSn M1 mice as previously described^68^ with the following modifications. Mice were anesthetized with isoflurane and CTX injection was performed on a single injection into the TA (50 μl, 10 μM) and muscle regeneration assessed after 96h. Muscle explants-derived EVs used in Fig. 1b-d were obtained from F2 cross between the offspring of *Myf5-Cre* mice^69^ and Wnt7a*^fl/fl^* mice^70^ in a C57BL/6 genetic background. Briefly, mice were anesthetized with isoflurane and CTX injection was performed on three injections (50 μl, 10 μM each), into the TA, and one on each lobule of the gastrocnemius. Hind limb muscles were harvested after 96h. *In vivo* assays in Fig 6 d-i were performed in 12-week-old male C57BL/10ScSn-Dmdmdx/J mice from The Jackson Laboratory, more details in the electroporation section.

### Pre-embedding immunogold labeling for tissue TEM

Briefly, TA specimens were fixed in Karnovskýs fixative for 2 weeks. After fixation all segments were subsequently washed with 0.1M sodium cacodylate, and treated with 0.1% sodium borohydride in PBS. Samples were permeabilized with 0.1% triton X-100 and blocked with 10% donkey serum + 0.6% fish gelatin. TA samples were incubated with Wnt7a antibody. After 48 h incubation, segments were rinsed thoroughly with PBS and incubated overnight with the secondary antibody Ultra small (0.8 nm) Gold conjugated (EMS) in blocking buffer at RT. Later, samples were rinsed with 0.1M sodium cacodylate and post-fixed with 2% glutaraldehyde in 0.1 M sodium cacodylate. Pre-embedding enhancement was realized with silver enhancement kit (AURION R-Gent SE-EM, EMS) according to the manufacturer’s instructions. After enhancement, samples were secondly postfixed with 1% osmium tetroxide in 0.1 M sodium cacodylate buffer. Then, samples were dehydrated in increasing concentration of ethanol and infiltrated in Spurr resin. Ultrathin transversal sections (80 nm) were collected onto 200-mesh copper grids and counterstained with 2% aqueous uranyl acetate and with Reynold’s lead citrate. Finally, specimens were observed under a transmission electron microscope (Hitachi 7100, Gatan digital camera). For our analysis, approximately 50 immunoelectron micrographs were examined per muscle at different magnifications.

### Pre-embedding immunogold labeling for cells and EVs TEM

Fixed HEK293T cells/EVs pellets were treated separately with 0.1% sodium borohydride in PBS. Half of the specimen pellets were permeabilized with 0.1% Triton X-100 for 10 minutes and the other half of the samples were processed without permeabilization. Pellets were blocked in blocking buffer (10% donkey serum + 0.6% gelatin from cold water fish skin in PBS) for 2 h. Pellets were incubated with the primary antibody for 48 h. Pellets were incubated overnight with the secondary antibody (Jackson ImmunoResearch). Immunogold-labelled cells and EVs were fixed with 2% glutaraldehyde in 0.1 M sodium cacodylate buffer and enhancement was performed with a silver enhancement kit on the immunogold-labelled cells. All samples were post-fixed with 1% osmium tetroxide in 0.1 M sodium cacodylate buffer. Specimens were dehydrated and embedded in resin and polymerized overnight at 70°C. Immunogold-labelled exosome ultrathin sections were observed by transmission electron microscopy at 100 000X and 150 000x.

### Conditioned media production for cell derived EVs

Equal numbers of HEK293T cells were seeded and the different plasmids were transfected with linear polyethylenimine (Polysciences), accordingly to manufacturer’s instructions. Later, transfected cells were cultured with 10% FBS exosome-depleted^67^ in DMEM (Gibco) and maintained at 37°C in a humidified incubator equilibrated with 5% CO2. After 48h of secretion conditioned media was collected for EVs isolation. For EVs derived from RPTEC hTERT1 cells, equal numbers of cells were seeded and let them grow in growth media, previously described. Upon 80% confluency growth media was change and maintained at 37°C in a humidified incubator equilibrated with 5% CO2 for 48h of secretion conditioned media. Then media was collected for EVs isolation following regular protocol.

### Conditioned media production for muscle explants derived EVs

For tissue EVs we have standardized a protocol to obtain conditioned media from muscles explants. Briefly, four days after injury both hind limbs were harvested and cultured as explants on an exosome-depleted FBS pre-coated dish with high-glucose DMEM (Gibco) and maintained at 37°C in a humidified incubator equilibrated with 5% CO2. After 48h conditioned media was collected for exosomal isolation.

### EVs isolation

Conditioned media (20mL) was clarified by sequential centrifugation (300g at RT for 10 min; 2500 g at RT for 10 min and 20,000 g at 4 °C for 20 min). Supernatant was transferred to Flexboy bag (Sartorius) and subjected to tangential flow filtration (TFF) under sterile conditions. Briefly, a KrosFlo Research 2i TFF system (Spectrum Laboratories) coupled to a MidGee Hoop ultrafiltration hollow fiber cartridge (GE Healthcare) 500-KDa MWCO was used. Transmembrane pressure was automatically adjusted at 3 PSI and a shear rate at 3000 s-1. Sample was concentrated up to 10mL and then subjected to continuous diafiltration. Finally, sample was concentrated at 5mL and recovered from the cartridge. Lastly, EVs were pellet down after spinning on an ultrabench centrifuge for 30min at 100,000 g at 4 °C.

### Immunoblot analysis

Immunoblot analysis was performed as described previously^71^ with the following modifications. The lysates from EVs were not clarified by centrifugation. The immunoblot transferring was performed onto PVDF membranes. All antibodies and dilutions are provided in Extended Data Table 3.

### Immunohistochemistry

TA muscle cryosections were rehydrated using PBS, and then fixed with 2% PFA in PBS at room temperature. After washing with PBS, permeabilization with a solution of 0.1% Triton and 0.1 M glycine in PBS was applied for 10 min at room temperature. Mouse on mouse blocking reagent was used at a dilution of 1:40 in blocking solution of 10% goat serum, 1% bovine serum albumin (BSA) and 0.1% Tween 20 in PBS for one hour at room temperature. Primary antibodies were incubated overnight. Nuclei were counterstained with DAPI before mounting in Permafluor. For analysis Z-stack images of cryosections were acquired on an epifluorescence microscope equipped with a motorized stage (Zeiss AxioObserver Z1) with a step size of 0.2 μm to span the cell (25 slices in total) and images were deconvoluted using Zen Software (Zeiss). 3D sum intensity Z-projection was performed with ImageJ software. Laminin staining was analyzed for fiber counting and minimum Feret’s diameter using SMASH^72^.

HEK293T transfected cells were fixed 10 min with 4% PFA, 10 min perm (0.1M glycine, 0.1% Tx100 in PBS), 1 hour blocking (5% HS, 2% BSA in PBS), o/n primaries at 4 degrees in blocking, and 1 hour secondary in blocking with washes in between. Hoechst was used for nuclei counterstain before mounting in Permafluor. Images were acquired using a Zeiss LSM900 with Airyscan and 63X oil objective.

### Hypertrophy assay

For Fig. 1c, d wild type primary myoblasts from C57BL/10ScSn M1 mice were differentiated for 4 days along with EVs stimulation at 10 μg/mL (based on total extracellular vesicle protein quantification after lysis) or recombinant Wnt7a protein at 100ng/mL.

For Fig. 6 b, c, Wnt7a primary myoblasts from C57BL/10ScSn M1 mice, previously obtained as aforementioned, were transfected twice (every 24h two consecutive days) with COPA, COPB2 and COPA+COPB2 siRNA at 10nM (final concentration) using Lipofectamine RNAiMax, accordingly to manufacturer’s instructions. After siRNA knockdown cells were differentiated for 3 days as aforementioned.

Upon three days of differentiation myotubes from both experiments were fixed with 4% PFA. Permeabilization and blocking solution consisting of 0.3 M glycine, 1% BSA and 0.1%Tween in PBS was added for 90 mins. p-MHC primary antibody was incubated overnight. Nuclei were counterstained with DAPI before mounting in Permafluor.

For analysis Z-stack images of myotubes were acquired on an epifluorescence microscope equipped with a motorized stage (Zeiss AxioObserver Z1) with a step size of 0.2 μm to span the cell (25 slices in total) and images were deconvoluted using Zen Software (Zeiss). 3D sum intensity Z-projection was performed with ImageJ software. Ten blind images were acquired per sample. The 50 largest myotubes from each well were included in the analysis. FIJI software was used to analyze myotube diameter. Three different samples were measured per condition.

### Construction of Wnt7a mutants

All constructs were cloned in a pcDNA3 vector. Wnt7a was originated from a pcDNA3-hWnt7a-HA plasmid that was designed in our lab. Wnt10a and Wnt16 were originated from pcDNA-hWnt10a-V5 (<u>Addgene</u> 35939) or pcDNA-hWnt16-V5 (<u>Addgene</u> 35942) plasmid respectively. Wnt10b and Wnt3a constructs used in this publication were a gift from Marian Waterman, David Virshup and Xi He from the plasmid kit^73^ (Addgene kit # 1000000022). HALO*EBP-HA and HALO*EBP constructs were originated from a pBSM13-Pax7HALO plasmid that was designed in our lab. Mutation and truncation were generated by overlap extension PCR with specially designed primers. BamHI and EcoRI restriction sites were included in primers. OE-PCR products and pcDNA3-HA vector were digested with BamHI and EcoRI and ligated with Takara ligase Solution. All constructs were verified by sequencing. All primers and coding sequences sources are provided in Extended Data Table 4.

### *In-silico* homology modeling of Wnt7a

The homology model of human Wnt7a was constructed through its sequence annealing over the resolved structure of Wnt3 protein (PDB^74^ 6AHY^75^) with FoldX^29^ BuildModel command. The annealing of the sequence resulted in no energetic conflicts enlighting that the folding captured by the crystal represents a stable configuration of proteins within the Wnt family. GSGS linker length was chosen to replace EBP. To affect folding, we considered the distance criteria respect to the terminal residues of the EBP.

### *In-silico* determination of the EBP region

The *in-silico* determination of the EBP region was performed through the free energy measurement of folding of the Wnt7a model (ΔG_wt_) versus the free energy resulting of the truncation of windows of 15 aa (ΔG_truncated_) along the whole sequence. Those regions not contributing to the protein folding present a very negative variation energy (ΔΔG_truncated_WT_<<0). The N-terminal region is not structured since the mapping of the Wnt folding domain (PFAM ^76^ PF00110) starts in position 41, the C-terminal region presents also low energies being a folded region not in close contact with the rest of the protein. Besides the terminal regions, the only sequence window presenting very low energy was selected as EBP (afterwards confirmed experimentally) since it is not important for the folding, is highly variable along the Wnt family, evincing that its sequence codifies for functional behavior.

### Modeling of the EBP-loop swapping

All the unstructured regions within the Wnt7a generated model that were surrounded by secondary structured regions were evaluated in terms of end-to-end distances and torsional angles to establish their ability to room the EBP region through a sequence swap. Using ModelX^77^ fragment replacement the EBP was inserted using as anchoring terminal aminoacids GLU171 and ASN 175. Energies of the replaced model was measured then with the FoldX force field and no energetic conflicts or clashes where found, demonstrating that the sequence swapping was supported by the structure.

### Uptake assays

HEK293T cells were transfected with pcDNA3_HALO and pcDNA3_HALO-EBP plasmids that we generated from a Pax7-HALO plasmid (Epoch Life Science) using PEI as aforementioned. EVs from transfected cells were isolated as previously described and added to fresh seeded HEK293T for 15 min. After, stimulated cells were labeled with HaloTag® Ligands for Super Resolution Microscopy-Janelia 549 (Promega) accordingly to manufacturer’s instructions. Cells were then fixed in 2% PFA for 5 min and washed three times with PBS. Lastly cells were analyzed by image cytometry in the Amnis ImageStream X platform to verify the location of the fluorescence within the cytoplasm. The fluorescence detected by the Amnis ImageStream was excited using 561nm laser and detected by the 580-30 emission filter channel.

### Inhibition of PORCN

HEK293T cells were transfected with pcDNA3_Wnt7a-FL-HA using PEI as aforementioned. After 6h of transfection cells were treated overnight with PORCN inhbibitor diluted in DMSO (AdooQ) at two different concentrations 100nM and 500nM in fresh media. Next morning media was changed for EVs secretion media and let them to secrete for 48h as aforementioned.

### BioID assay

Stable primary myoblast cell lines expressing BioID2, BioID2-EBP, or Wnt7a-BioID2 fusion proteins were generated using the mycBioID2-pBABE-puro vector (Addgene Plasmid). Myoblasts were grown in 15 cm culture dishes at sub confluency and incubated with biotin (Sigma-Aldrich: dissolved in DMSO) at a final concentration of 50 uM for 18 h. Plates were scraped in ice cold PBS, spun at 20817 g for 5 min to concentrate cell pellet, then resuspended in RIPA lysis buffer containing protease inhibitor cocktail. Cells were incubated on ice for 30 min, and then spun down 20817 g at 4°C for 20 min. Supernatant was transferred to new low retention Eppendorf tube, and protein concentration was quantified using Bradford reagent and spectrometry. Magnetic streptavidin beads (New England Biolabs) were used to precipitate the biotinylated protein fraction. Streptavidin beads were washed twice in RIPA lysis buffer and subsequently added to protein lysates for overnight incubation at 4°C rotating. The following day, beads were sequentially washed with RIPA buffer, 1 M KCl, 0.1 M Na_2_CO_3_, 2 M urea in 10 mM Tris-HCl (pH 8), and a final RIPA buffer wash. Biotinylated proteins were then eluted from beads by boiling for 10 min in 25 ul 6x Laemmli buffer containing 20 mM DTT and 2 mM biotin. Supernatant was loaded into precast gradient gel (4-15% Mini-PROTEAN® TGX Stain-Free™ Protein Gel) and run for 30 min at 100V. Colloidal blue dye (Thermofisher:) was applied for 3 h, then rinsed in Mili-Q water while shaking overnight. The entire protein containing lane for each condition was then cut out and stored in 1% acetic acid. Samples were then transferred to the Ottawa Hospital Research Institute Proteomics Core Facility for further processing as described below.

### Proteomic analysis

Proteins were digested in-gel using trypsin (Promega) according to the method of Shevchenko^78^. Peptide extracts were concentrated by Vacufuge (Eppendorf) and purified by ZipTip (Sigma-Millipore). LC-MS/MS was performed using a Dionex Ultimate 3000 RLSC nano HPLC (Thermo Scientific) and Orbitrap Fusion Lumos mass spectrometer (Thermo Scientific). MASCOT software version 2.6.2 (Matrix Science) was used to infer peptide and protein identities from the mass spectra. The observed spectra were matched against sequences from SwissProt (version 2020-01) and against an in-house database of common contaminants. The results were exported to Scaffold (Proteome Software) for further validation and viewing. Enrichment heatmap was generated by computing the log_2_ of the fold enrichment of each condition versus its control. Gene Ontology term enrichment analysis was performed over the “cellular component” branch using ClueGO plugin on Cytoscape software.

### Proximity ligation assay (PLA)

Fixed cells (Murine primary myotubes, RPTEC hTERT or transfected HEK293T cells) were permeabilized (0.1% Triton X-100, 0.1 M Glycine, PBS) for 10 min and blocked with Duolink Blocking Solution (Sigma) for 3h. Incubation with primary antibodies diluted in Duolink Blocking Solution (Sigma) was performed overnight at 4°C. PLA reactions were subsequently performed using Duolink PLA probes for goat-mouse and goat-rabbit and Duolink *In Situ* Detection Reagents Red (Sigma) following the manufacturer’s protocol. Myotubes were counterstained with GM130 to visualize the Golgi Apparatus. After the final wash, cells were mounted with VECTASHIELD Antifade Mounting Medium with DAPI (Vector Laboratories). For analysis Z-stack images of myotubes were acquired on an epifluorescence microscope equipped with a motorized stage (Zeiss AxioObserver Z1) with a step size of 0.2 μm to span the cell (25 slices in total) and images were deconvoluted using Zen Software (Zeiss). 3D sum intensity Z-projection was performed with ImageJ software.

### Immunoprecipitation

Wnt7a-HA was overexpressed in HEK293T cells using Lipofectamine 2000 (Life Technologies) according to the manufactureŕs instructions. Cells and EVs were isolated two days post-transfection and lysed in immunoprecipitation lysis buffer (50 mM Tris pH 7.5, 150 mM NaCl, 2 mM MgCl2, 0.5 mM EDTA, 0.5% Triton X-100, and protease inhibitors) for 30 min on ice. Lysates from cells were cleared by centrifugation and were incubated with either HA (Benthyl) or COPB2 (Cusabio) antibodies-Dynabeads Protein G (Thermo Fisher) overnight at 4°C, accordingly to the manufacturer’s instructions. Beads were washed 4 times with lysis buffer and eluted with Laemmli buffer. Immunoprecipitates were resolved by SDS-PAGE and analyzed by immunoblot with the indicated antibodies.

### SiRNA silencing

siRNA transfections were performed on HEK293T cells at 16 h post-culture using Lipofectamine RNAiMAX (Life Technologies) according to the manufactureŕs instructions. siRNAs for WLS, COPA, COPB2 were purchased from Dharmacon and used at a final concentration of 10nM, 10nM and 20 nM respectively. The following day cells were transfected with Wnt7a as aforementioned and Wnt7a secretion was tested 48h later.

### Recombinant protein production

The DNA sequence encoding COPB2_1-304_ was custom synthesized with codon optimization for expression in E. coli (GenScript) and cloned into pET28-Sumo3 vector (EMBL, Heidelberg) to express it with a N-terminal cleavable 6xHis-Sumo3 tag.

### Protein expression and purification

Native protein was expressed in *E. coli* BL21(DE3) grown in Luria-Bertani (LB) broth at 37 °C, and protein expression was induced at an OD600 of 0.8 by the addition of 0.5 mM isopropyl-β-D-thiogalactopyranoside (IPTG). Cells were harvested after 16 hours of growth at 18 °C. All following purification steps were performed at 4 °C. The concentration of all purified proteins was calculated using the theoretical extinction coefficient.

For the purification of COPB2_1-304_, cell pellets were lysed at 4°C using high-pressure homogenization at 25 Kpsi (1 Kpsi = 69 _ 103 MPa) (Constant System Ltd, UK) in lysis buffer [50 mM Tris-HCl pH:7.5, 500 mM NaCl, 1 mM dithiothreitol (DTT), 10 mM imidazole] and supplemented with 0.1 mM phenylmethylsulfonyl fluoride (PMSF) and 1 mM benzamidine. After centrifugation at 50,000 x g for 45 min, the soluble fraction was incubated for 2 hours in batch with Ni2+-nitrilotriacetate (NTA) agarose resin (Macherey-Nagel). After extensive washing of the beads with lysis buffer, the protein was eluted with lysis buffer supplemented with 250 mM imidazol. The N-terminal 6xHis-Sumo3-tag was subjected to overnight Sentrin-specific protease 2 (SENP2) digestion via dialysis into 150 mM NaCl, 1 mM DTT, 10 mM imidazole and 25 mM Tris-HCl pH 7.5. A second Ni2+-NTA chromatography was carried out to remove the cleaved tag and uncleaved protein. COPB2_1-304_ was subsequently purified by ion-exchange chromatography (HitrapQ, GE Healthcare), diluting the NaCl concentration to 30mM with buffer A (25mM Tris-HCl 7.0, 1mM DTT) and using a gradient of 30–1000 mM NaCl followed by size-exclusion chromatography (Superdex 75 16/60, GE Healthcare) in buffer B [25 mM Tris-HCl pH 7.0, 150 mM NaCl, 0.5 mM Tris(2-carboxyethyl) phosphine (TCEP)].

### Crystallization, data collection

COPB2_1-304_ at 10mg/ml (0.28mM) was incubated with the Wnt_252-257_ peptide (LKIKKP) at 3mM during one hour in 150mM NaCl, 25mM Tris pH 7.5 before setting the crystallization experiments. Initial screening was performed with a Mosquito robot (TTP, Labtech) in 96-well MRC plates (Molecular Dimensions) at 18 °C. A total of 576 different crystallization conditions from commercial screenings (Molecular Dimensions, Hampton Research & Quiagen) were tested. The sitting drop experiments were performed with 250 nL of protein/peptide plus 250 nL from the reservoir solution. Crystal hits were manually optimized with drops containing 2 μL of protein and 2 μL of reservoir solution. Best crystals were obtained in 0.1M Hepes pH 7.0 and 2.0M AmSO_4_. Crystals were cryoprotected with 20% glycerol, and diffraction data were collected in the XALOC beamline at ALBA synchrotron (Extended Data Table 2). Diffraction data were indexed, integrated and scaled in P1 using XDS^46^. PHASER^79^ was used to solve the structure by molecular replacement using COPB2_1-304_ (PDB 2YNN) as a search model. The final model was obtained after several cycles of manual modification using COOT^80^ followed by refinement using 6 twin operators in REFMAC^81^. The coordinates and structure factors have been deposited at the Protein Data Bank (PDB) with the entry code PDB: 8A86.

### Isothermal titration calorimetry assays

Isothermal titration calorimetry (ITC) experiments were carried out on a VP-ITC titration microcalorimeter (MicroCal/GE Healthcare) at 25 °C. COPB2_1-304_ and peptides used for ITC experiments were dialyzed overnight at 4 °C against 150 mM NaC, 0.5 mM TCEP and 25 mM HEPES 7.5, and degassed for 5 minutes in a ThermoVac sample degasser before titration. The titration sequence consisted of an initial 2 μl injection to prevent artifacts arising from filling of the syringe (not used in data fitting), followed by 28 injections of 10 μl aliquots with a spacing of 210 s between injections. Similar injections of protein or peptides in buffer were performed to determine the heat of dilution used to correct the experimental data. The resulting titration data were integrated and fitted to a one-site model using the Origin ITC software package supplied by MicroCal. The binding constant (Ka, Kd=1/Ka), the molar binding stoichiometry (n) and binding enthalpy (ΔH) were extracted directly from the fit. The free energy (ΔG) and entropy (ΔS) of binding was calculated from ΔG = −RTlnKa = ΔH-TΔS, where R is the gas constant and T is the absolute temperature. The interaction of Wnt peptides with COPB2_1-304_ were analyzed by titrating 450 μM of each peptide into 20 μM COPB2_1-304_. Data are the mean of a minimum of three replicate titrations for each experiment.

### Electroporation

Right TA muscle was electroporated either with 30ug of pcDNA3_Wnt7a or pcDNA3_ Wnt7a_ΔEBP*GSGS. Contralateral legs were electroporated with empty vector pcDNA3. All plasmids were purified using endotoxin-free kits (Quiagen) and diluted in saline. 50uL of plasmid were injected intramuscularly into the TA of mdx mice under general anesthesia. Electric stimulation was applied with a pulse generator (ECM 830, BTX) of 100-150 volts for 6 pulses, with a fixed duration of 20ms and an interval of 200ms using 5mm needle electrodes (BTX).

### Statistical analysis

Experiments were performed with a minimum of three biological replicates and results are presented as the mean ± SEM. Student’s t-test were performed to assess the statistical significance of two-tailed analysis. For multiple comparisons ANOVA test was employed and TUKEY test for post-hoc analysis. *P*-values are indicated as *p ≤ 0.05, **p ≤ 0.01, ***p ≤ 0.001, and *P-*values <0.05 were considered statistically significant. The exact *p* values are provided in the Source Data file.

## Supporting information

Extended DataTable 2

## Acknowledgements

The authors thank Jennifer Richie for the mice colony management, Lawrence Puente for mass spectrometry, Fernando Ortiz for flow cytometry, Adam Smith for particle isolation, Dylan Burger for his advice on extracellular vesicle characterization and the NTA equipment, the Ottawa Bioinformatics Core Facility, Claire Lastrucci for her help with graphical conceptualization, and Vivek Malhotra for critical reading the manuscript. D.D. holds a Frederick Banting and Charles Best Canada Graduate Scholarships - Doctoral Award (CGS-D). This study made use of the ALBA synchrotron beamline BL13-XALOC with the collaboration of ALBA staff. The studies from the laboratory of M.A.R. were carried out with support of grants from Defeat Duchenne Canada, the US National Institutes for Health [R01AR044031], the Canadian Institutes for Health Research [FDN-148387], Ontario Institute for Regenerative Medicine, and the Stem Cell Network.

## Author contributions

U.G.R: Conceptualization, Methodology, Validation, Formal Analysis, Investigation, Data Curation, Writing-Original draft, Visualization, Supervision and Funding acquisition; D.D: Conceptualization, Methodology, Validation, Investigation, Data Curation, Writing-Review & Editing; L.R: Conceptualization, Methodology, Software, Formal analysis, Data Curation, Writing-Review & Editing; M.E.: Investigation; S.F: Investigation; F.X: Investigation, Resources; H.M: Investigation; M.A: Investigation; Y.D.R: Investigation; R.K: Writing-Review & Editing; A.L.R: Methodology, Validation, Software, Data Curation, formal analysis; L.S: Conceptualization, Writing-Review & Editing; A.H: Conceptualization, Formal analysis, Data Curation, Writing-Review & Editing, M.A.R.: Conceptualization, Formal Analysis, Data Curation, Writing-Original draft, Writing-Review & Editing, Supervision, Project administration and Funding acquisition.

## EXTENDED DATA FIGURE LEGENDS

**Extended Data Fig. 1 Wnt7a is secreted in several modes and Tangential Flow Filtration allows separation of the different types of secretion**. a, iTEM of anti-HA labeling of Wnt7a-HA transfected HEK293T cells shows both types of Wnt7a secretion, on exosomes surface (arrowheads) and as free protein (arrows). Scale bar 100nm. b, Representative images of anti-Lamp1 and anti-M6PR immunofluorescence in HEK293T cells showing no merge with Wnt7a. Cells were counterstained with DAPI. Scale bar 5μm. c, Immunoblot analysis of Wnt7a-EVs derived from HEK293T that are retained inside the TFF cartridge within the retentate fraction, and free-Wnt7a passes through the pores of the column and is collected in the permeate fraction. Wnt7a co-purified with EVs together with the exosomal protein CD81. d, Quantification of Wnt7a expression on secreted EVs surface versus free protein secretion derived from HEK293T. e, Relative size distribution analysis of EVs fraction from HEK293T cells comparing EVs derived from Empty Vector transfected HEK293T cells versus Wnt7a transfected HEK293T cells. f, iTEM of anti-HA labeling of EVs from HEK293T Wnt7a-HA transfected cells, showing HA expression on EVs surface. n = 3 biological replicates. Data are mean ± s.e.m.

**Extended Data Fig. 2 Tangential Flow Filtration methodology for isolation of exosomes from muscle explants**. **a**, Experimental protocol used to obtain EVs from mice hind limb muscle. **b**, Relative size distribution analysis of EVs fraction from muscle explants. **c**, Immunoblot analysis of EVs fraction from muscle showing Wnt7a expression. **d**, Hypertrophy dose-response assay of murine primary myotubes treated with muscle EVs. Data shown as fold change on myotube diameter over the control (%); Wnt7a recombinant protein was used as a positive control. **e**, pMHC immunofluorescence representative images of hypertrophied myotubes after muscle EVs stimulation. Scale bar 50μm. **f**, Immunofluorescence confirmation of Wnt7a expression abrogation in *Myf5*(Cre/+)*: Wnt7a*(fl/fl) injured TA at 96h post-CTX injury. Scale bar 50μm. **g**, Immunoblot verification of Wnt7a expression abrogation in EVs isolated *from Myf5*(Cre/+)*: Wnt7a*(fl/fl) hind limb muscle at 96 h post-CTX injury. **h,** iTEM of anti-Wnt7a labeling of EVs showing abrogation of Wnt7a expression in EVs from *Myf5*(Cre/+)*: Wnt7a*(fl/fl) mice muscle explants. Scale bar 100 nm. n = 3 biological replicates. Data are mean ± s.e.m.

**Extended Data Fig. 3 Wnt7a secretion is regulated by the Exosomes Binding Peptide. a**, Graph display quantitative secretion analysis of each truncate from Fig. 2b. Data shown as the ratio between EVs and cells fractions. (n=3, Data are mean ± s.e.m., ANOVA test p-value 2.62E-06, TUKEY test **≤0.01, ***≤0.001). **b**, Wnt7a protein tertiary structure highlighting the Wnt7a minimal structure required for EV secretion.

**Extended Data Fig. 4 The EBP targets proteins for extracellular secretion on exosomes. a**, 3D modeling of EBP insertion in a similar structural space. (**a**-Left lower) In blue the EBP and the replaced region (AAs 172-174), in red the aminoacids anchoring both unstructured regions. The small difference in Cα-Cα distance of residues anchoring both peptides gives room to swap them, considering as well that are in the same face of the structural surface. (**a**-Right upper) In green, the EBP modeled into the replaced region (AAs 172-174), side chains in sticks. **b**, Representative images of HEK293T producing cells upon incubation with HALO fluorescent tag, proving the overexpression of HALO and HALO-EBP protein. **c**, Scheme of the protocol to visualize uptake of HALO*EBP EVs by Image Cytometry.

**Extended Data Fig. 5 Wnt7a secretion on exosomes is mediated by interaction with Coatomer proteins. a**, Immunoblot EVs secretion analysis in HEK293T cells of the minimal Wnt7a structure necessary for EVs secretion from 100-300 aa (right panel) with (Wnt7a_Δ32-99 Δ301-349) and without Signal Peptide (Wnt7a_Δ1-99 Δ301-349). Signal peptide is not required for EVs-Wnt7a secretion. **b,** Quantification of Wnt7a Δ301-349 secretion comparing with (Wnt7a_Δ32-99 Δ301-349) and without Signal Peptide (Wnt7a_Δ1-99 Δ301-349) in HEK293T cells. **c,** Immunoblot secretion analysis in HEK293T cells confirms secretion of the different Wnt7a truncates from Figure 2a without the Signal Peptide. **d,** qPCR analysis of WLS expression upon siRNA WLS in HEK293T cells. **e,** Experimental scheme of the BirA assay protocol. **f**, Immunoblot analysis of BioID constructs from primary myoblasts expressing WT, BirA-myc tagged, Wnt7a-BirA-myc tagged and EBP-BirA-myc tagged. **g,** Quantification of Wnt7a mass spectrometry counts for EVs derived from primary myoblasts expressing Wnt7a-BirA-myc tagged, EBP-BirA-myc tagged, and BirA-myc tagged. **h-i,** iTEM representative images for anti-Wnt7a and anti-Myc labeling of EVs derived from primary myoblasts expressing Wnt7a-BirA-myc tagged, EBP-BirA-myc tagged, and BirA-myc tagged with and without permeabilization.

**Extended Data Fig. 6 Wnt7a secretion on exosomes is mediated by interaction with Coatomer proteins. a,** Gene Ontology (GO) term fold enrichment analysis for the gene set displayed in Fig. 4a. The graph displays terms along the hierarchy within the “cellular component” branch, the analysis was performed using ClueGO plugin on Cytoscape software. **b,** Wnt7a:COPB2 PLA (red) performed in murine primary myotubes either expressing Wnt7a-BirA or BirA. PLA signal was counterstained with GM310 (green), a Golgi Apparatus marker and with DAPI (blue), showing interaction in the plasma membrane area. Scale bar 10 μm. **c,** qPCR analysis of Wnt7a endogenous expression on RPTEC-hTERT cells. HEK293T cells were used as negative control and GAPDH, Ppia and RPS18 as housekeeping genes. **d,** Representative immunofluorescence images of Wnt7a endogenous expression on RPTEC-hTERT cells. Counterstaining was done with WGA, a plasma membrane marker and DAPI. No Wnt7a antibody added was used as a negative control. **e,** Representative immunofluorescence images of Wnt7a transfected truncates in HEK293T cells. **f,** Quantification of total mass spectrometry counts for Wnt7a-HA, COPA, COPB2 upon HA immunoprecipitation in EVs derived from Wnt7a-HA transfected HEK293T cells.

**Extended Data Fig. 7 The KIK motif within the EBP regulates binding to the Coatomer complex. a**, Overlap of the KTKLL motif of Emp47p (colored blue) and the KTKKP motif of Wnt7a (orange) bound to the COPB2 N-terminal WD-repeat domain. The average root means square deviation (RMSD) for the all-atom pairwise superposition was 0.33 Å. The picture is meant to emphasize that the two KxKxx motifs adopt the same conformation.

**Extended Data Fig. 8 Exosomal Wnt secretion regulated by EBP is a conserved mechanism. a**, Alignment of Wnt family proteins showing in green the conservation degree of the KKx, RR, KHR, RxR, KR, RR positively charged motifs among the Wnt family. **b**, Homology model of Wg showing in orange the EBP region and in the left panel the EBP sequence containing the KR and KK motifs in cyan. **c**, Alignment of EBP-Wnt7a sequences showing the conservation degree across the species.

**Extended Data Fig. 9 EBP sequence is required for a comprehensive regenerative muscle response mediated by Wnt7a-Coatomer interaction**. a, qPCR analysis of COPA, COPB2 and Wnt7a expression upon siRNA COPA, COPB2 in mouse primary myoblasts. b, Representative immunofluorescence images of the entire TA muscle cryosection 21 days after electroporation of Wnt7a or Wnt7a_ΔEBP*GSGS plasmids. c, Representative H&E images of the entire TA muscle cryosection 21 days after electroporation of Wnt7a or Wnt7a_ΔEBP*GSGS plasmids.

