## Extended Data Table 1 for "Wnt binding to Coatomer proteins directs secretion on exosomes independently of palmitoylation"

**Extended Data Table 1.** Thermodynamic data from the isothermal titration calorimetry study to characterize the interaction between EBP and COPB2

| Injectant | Cell | Kd (μM) | n | ΔH (kcal/mol) | -TΔS (kcal/mol) | ΔG (kcal/mol) |
|---|---|---|---|---|---|---|
| EBP(aa240-257)PVRASRNKRPTFLKIKKP | COPB2(1-304) | 11,61 | 0,97 | -13.33±0.71 | 6.59 | -6,74 |
| EBP C-terminal half (aa252-257) LKIKKP | COPB2(1-304) | 37,17 | 0,82 | -19.07±2.99 | 13.02 | -6,05 |
| EBP N-terminal half (aa244-250) SRNKRPT | COPB2(1-304) | N.B. | | | | |
| EBP(K253A) PVRASRNKRPTFLAIKKP | COPB2(1-304) | 47,17 | 1,18 | -6.59±1.20 | 0.69 | -5,9 |
| EBP(K255A) PVRASRNKRPTFLKIAKP | COPB2(1-304) | N.B. | | | | |
| EBP(K256A)PVRASRNKRPTFLKIKAP | COPB2(1-304) | 5,99 | 0,95 | -7.63±0.30 | 0.50 | -7,13 |
