## Extended DataTable 2 for "Wnt binding to Coatomer proteins directs secretion on exosomes independently of palmitoylation"

**Extended Data Table 2.** Data collection and refinement statistics for crystallography.

| | COPB2 $_{1-304}$ / Wnt7a $_{252-257}$ |
|---|---|
| **Data collection** | |
| Wavelength [Å] | 0.9793 |
| Space group | P1 |
| Resolution [Å] | 50.0-1.81 (1.92-1.81) |
| Cell dimensions | |
| *a, b, c* [Å] | 1.5, 159.5, 159.6 |
| α, β, γ [º] | 107.3, 107.3, 107.4 |
| $CC_{1/2}$ (%) | 99.6(31.0) |
| Completeness (%) | 95.5 (91.3) |
| I/σ | 8.53 (0.72) |
| Number of unique reflexions | 1157927(178778) |
| Redundancy | 3.5 (3.6) |
| **Refinement** | |
| R-factor (%) | 15.9 |
| R-free (%) | 17.4 |
| No. atoms: | |
| Waters | 3348 |
| Ions/$SO_4$ | 206 |
| Ligand/Glycerol | 57 |
| R.m.s deviations: | |
| Bond lengths (Å) | 0.002 |
| Bond angles (º) | 1.24 |
| **PDB CODE** | **8A86** |

*Highest resolution shell is shown in parenthesis.
