## Extended Data Table 3 for "Wnt binding to Coatomer proteins directs secretion on exosomes independently of palmitoylation"

### Extended Data Table 3. Antibodies used in this study.

| Antibody | Application | Size | Species | Dilution | Source | Secondary Antibody |
|---|---|---|---|---|---|---|
| **Calnexin** | Immunoblot | 90KDa | Rabbit | 1:5000 | Abcam, ab22595 | Anti-rabbit 1:5000 |
| **CD81** | Immunoblot | 18KDa | Rabbit | 1:500 | Saint Johns Lab, STJ96759 | Anti-rabbit 1:5000 |
| **CD9** | Immunoblot | 25KDa | Rabbit | 1:1000 | SBI EXOAB-CD9A-1 | Anti-rabbit 1:5000 |
| **COPα** | Proximity Ligation Assay | | Mouse | 1:50 | Santa Cruz Biotechnology sc-398099 | |
| **COPα** | Immunoblot | 140KDa | Mouse | 1:500 | Santa Cruz Biotechnology sc-39809 | Anti-mouse 1:5000 |
| **COPβ2** | Proximity Ligation Assay | | Mouse | 1:20 | Nobus NB600-102 | |
| **COPβ2** | Immunoblot | 103KDa | Rabbit | 1:1000 | Cusabio PA529993ESR1HU-100UL | Anti-rabbit 1:5000 |
| **GAPDH** | Immunoblot | 37KDa | Goat | 1:1000 | Sigma-Aldrich PLA0302-100UL | Anti-goat 1:5000 |
| **GM130** | Proximity Ligation Assay | | Rabbit | 1:100 | Abcam ab52649 | |
| **HA** | iTEM cells | | Rabbit | 1:20 | Bethyl, A190-108A | 6 nm gold Donkey anti Rabbit. 1:50 |
| **HA** | Immunoblot | 1KDa | Rabbit | 1:1000 | Bethyl, A190-108A | Anti-rabbit 1:5000 |
| **HA** | iTEM Evs | | Rabbit | 1:20 | Bethyl, A190-108A | 12 nm gold Donkey anti Rabbit. 1:50 |
| **HALO** | Fluorescence | | | 200 nM | Promega GA1110 | |
| **HALO** | Immunoblot | 33KDa | Mouse | 1:1000 | Promega G9211 | Anti-mouse 1:5000 |
| **HSP70** | Immunoblot | 70KDa | Rabbit | 1:1000 | SBI EXOAB-Hsp70A-1 | Anti-rabbit 1:5000 |
| **Laminin** | Immunofluorescence | | Rabbit | 1:1000 | Sigma L9393 | Alexa Fluor 546 (Invitrogen) 1:1000 |
| **Lamp2** | Immunofluorescence | | Rabbit | 1:200 | Abcam ab18528 | Alexa Fluor 546 (Invitrogen) 1:1000 |
| **M6PR-AF555** | Immunofluorescence | | Rabbit | 1:100 | Abcam ab203438 | Recombinant antibody with Alexa Fluor 555 |
| **Myc** | Immunoblot | 57KDa | Rabbit | 1:1000 | Bethyl A190-105A | Anti-rabbit 1:5000 |
| **pMyosin** | Immunofluorescence | | Mouse | 1:20 | Hybridoma Bank MF20 | Alexa Fluor 488 (Invitrogen) 1:1000 |
| **Wnt7a** | iTEM muscle | | Goat | 50ug/mL | R&D Systems AF3008 | 0.8nm gold Donkey anti Goat. 1:50 |
| **Wnt7a** | iTEM Evs | | Goat | 20ug/mL | R&D Systems AF3008 | 12nm gold Donkey anti Goat. 1:50 |
| **Wnt7a** | Immunofluorescence | | Goat | 10ug/mL | R&D Systems AF3008 | Alexa Fluor 488 (Invitrogen) 1:1000 |
| **Wnt7a** | Immunoblot | 39KDa | Goat | 1:2000 | R&D Systems AF3008 | Anti-goat 1:5000 |
| **Wnt7a** | Proximity Ligation Assay | | Goat | 10ug/mL | R&D Systems AF3008 | |
