## Extended Data Table 4 for "Wnt binding to Coatomer proteins directs secretion on exosomes independently of palmitoylation"

### Extended Data Table 2. Primers used in this study.

| Clone | Amino acid sequences | Forward primer | Reverse primer | Epitope Tags |
|---|---|---|---|---|
| **Wnt7a_Δ32-212** | MNRKARRCLGHLFLSLGMVYLRIGGFSSVVATCWTTLPQFR<br>ELGYVLKDKYNEAVHVE**PVRASRNKRPTFLKIKKP**LSYRKP<br>MDTDLVYIEKSPNYCEEDPVTGSVGTQGRACNKTAPQASG | gcttctcctcagtggtagctacgtgctgg<br>accacactgccac | aagaattctcaagcgtaatctggaac<br>atcgtatgggtacttgcacgtgtacat<br>ctccg | HA |
| **Wnt7a_Δ32-149** | MNRKARRCLGHLFLSLGMVYLRIGGFSSVVAGGCSADIRYG<br>IGFAKVFVDAREIKQNARTLMNLHNNEAGRKILEENMKLECK<br>CHGVSGSCTTKTCWTTLPQFRELGYVLKDKYNEAVHVE**PVR**<br>**ASRNKRPTFLKIKKP**LSYRKPMDTDLVYIEKSPNYCEEDPVT | aaggatccaccatgaaccggaaagcg<br>cggcgctgcctgggccacctctttctca<br>gcctgggcatggtctacctccggatcgg<br>tggcttctcctcagtggtagctggtggct | aagaattctcaagcgtaatctggaac<br>atcgtatgggtacttgcacgtgtacat<br>ctccg | HA |
| **Wnt7a_Δ32-99** | MNRKARRCLGHLFLSLGMVYLRIGGFSSVVAGSREAAFTYAI<br>IAAGVAHAITAACTQGNLSDCGCDKEKQGQYHRDEGWKWG<br>GCSADIRYGIGFAKVFVDAREIKQNARTLMNLHNNEAGRKIL<br>EENMKLECKCHGVSGSCTTKTCWTTLPQFRELGYVLKDKYN<br>EAVHVE**PVRASRNKRPTFLKIKKP**LSYRKPMDTDLVYIEKSP | aaggatccaccatgaaccggaaagcg<br>cggcgctgcctgggccacctctttctca<br>gcctgggcatggtctacctccggatcgg<br>tggcttctcctcagtggtagctgggagc<br>[illegible] | aagaattctcaagcgtaatctggaac<br>atcgtatgggtacttgcacgtgtacat<br>ctccg | HA |
| **Wnt7a_Δ32-49** | MNRKARRCLGHLFLSLGMVYLRIGGFSSVVAAICQSRPDAII<br>VIGEGSQMGLDECQFQFRNGRWNCSALGERTVFGKELKVG<br>SREAAFTYAIIAAGVAHAITAACTQGNLSDCGCDKEKQGQY<br>HRDEGWKWGGCSADIRYGIGFAKVFVDAREIKQNARTLMNL<br>HNNEAGRKILEENMKLECKCHGVSGSCTTKTCWTTLPQFRE | aaggatccaccatgaaccggaaagcg<br>cggcgctgcctgggccacctctttctca<br>gcctgggcatggtctacctccggatcgg<br>tggcttctcctcagtggtagctgcgatct<br>gccagagccggcccgac | aagaattctcaagcgtaatctggaac<br>atcgtatgggtacttgcacgtgtacat<br>ctccg | HA |
| **Wnt7a_Δ213-349** | MNRKARRCLGHLFLSLGMVYLRIGGFSSVVALGASIICNKIP<br>GLAPRQRAICQSRPDAIIVIGEGSQMGLDECQFQFRNGRWN<br>CSALGERTVFGKELKVGSREAAFTYAIIAAGVAHAITAACTQ<br>GNLSDCGCDKEKQGQYHRDEGWKWGGCSADIRYGIGFAK | aaggatccaccatgaaccggaaagcg<br>cggcgctg | aagaattctcaagcgtaatctggaac<br>atcgtatgggtacttggtggtgcacg<br>agcctgac | HA |
| **Wnt7a_Δ251-349** | MNRKARRCLGHLFLSLGMVYLRIGGFSSVVALGASIICNKIP<br>GLAPRQRAICQSRPDAIIVIGEGSQMGLDECQFQFRNGRWN<br>CSALGERTVFGKELKVGSREAAFTYAIIAAGVAHAITAACTQ<br>GNLSDCGCDKEKQGQYHRDEGWKWGGCSADIRYGIGFAK<br>VFVDAREIKQNARTLMNLHNNEAGRKILEENMKLECKCHGV | aaggatccaccatgaaccggaaagcg<br>cggcgctg | aagaattctcaagcgtaatctggaac<br>atcgtatgggta<br>ggtgggccgcttgttgcggctg | HA |
| **Wnt7a_Δ301-349** | MNRKARRCLGHLFLSLGMVYLRIGGFSSVVALGASIICNKIP<br>GLAPRQRAICQSRPDAIIVIGEGSQMGLDECQFQFRNGRWN<br>CSALGERTVFGKELKVGSREAAFTYAIIAAGVAHAITAACTQ<br>GNLSDCGCDKEKQGQYHRDEGWKWGGCSADIRYGIGFAK<br>VFVDAREIKQNARTLMNLHNNEAGRKILEENMKLECKCHGV | aaggatccaccatgaaccggaaagcg<br>cggcgctg | aagaattctcaagcgtaatctggaac<br>atcgtatgggtactggggagccgtct<br>tgttgcag | HA |
| **Wnt7a_Δ32-99_Δ301-349** | MNRKARRCLGHLFLSLGMVYLRIGGFSSVVAGSREAAFTYAI<br>IAAGVAHAITAACTQGNLSDCGCDKEKQGQYHRDEGWKWG<br>GCSADIRYGIGFAKVFVDAREIKQNARTLMNLHNNEAGRKIL<br>EENMKLECKCHGVSGSCTTKTCWTTLPQFRELGYVLKDKYN | aaggatccaccatgaaccggaaagcg<br>cggcgctgcctgggccacctctttctca<br>gcctgggcatggtctacctccggatcgg | aagaattctcaagcgtaatctggaac<br>atcgtatgggtactggggagccgtct<br>tgttgcag | HA |
| **Wnt7a_Δ1-99_Δ301-349** | GSREAAFTYAIIAAGVAHAITAACTQGNLSDCGCDKEKQGQ<br>YHRDEGWKWGGCSADIRYGIGFAKVFVDAREIKQNARTLMN<br>LHNNEAGRKILEENMKLECKCHGVSGSCTTKTCWTTLPQFR<br>ELGYVLKDKYNEAVHVE**PVRASRNKRPTFLKIKKP**LSYRKP | aaggatccaccatggggagccgggag<br>gctgcgttc | aagaattctcaagcgtaatctggaac<br>atcgtatgggtactggggagccgtct<br>tgttgcag | HA |

| | | | | |
|---|---|---|---|---|
| **Wnt7a_ΔEBP*GSGS** | MNRKARRCLGHLFLSLGMVYLRIGGFSSVVALGASIICNKIP GLAPRQRAICQSRPDAIIVIGEGSQMGLDECQFQFRNGRWN CSALGERTVFGKELKVGSREAAFTYAIIAAGVAHAITAACTQ GNLSDCGCDKEKQGQYHRDEGWKWGGCSADIRYGIGFAK VFVDAREIKQNARTLMNLHNNEAGRKILEENMKLECKCHGV | vvcaacgaggccgttcacgtggagcctggttcaggttcactgtcgtaccgcaagcccatggacacggac | gtccgtgtccatgggcttgcggtacgacagtgaacctgaaccaggctccacgtgaacggcctcgttg | HA |
| **Wnt7a_Δ1-49** | AICQSRPDAIIVIGEGSQMGLDECQFQFRNGRWNCSALGER TVFGKELKVGSREAAFTYAIIAAGVAHAITAACTQGNLSDCG CDKEKQGQYHRDEGWKWGGCSADIRYGIGFAKVFVDAREI KQNARTLMNLHNNEAGRKILEENMKLECKCHGVSGSCTTKT | aaggatccaccatggcgatctgccagagccggcccgac | aagaattctcaagcgtaatctggaacatcgtatgggtacttgcacgtgtacatctccg | HA |
| **Wnt7a_Δ1-99** | GSREAAFTYAIIAAGVAHAITAACTQGNLSDCGCDKEKQGQ YHRDEGWKWGGCSADIRYGIGFAKVFVDAREIKQNARTLMN LHNNEAGRKILEENMKLECKCHGVSGSCTTKTCWTTLPQFR ELGYVLKDKYNEAVHVE**PVRASRNKRPTFLKIKKP**LSYRKP | aaggatccaccatggggagccgggaggctgcgttc | aagaattctcaagcgtaatctggaacatcgtatgggtacttgcacgtgtacatctccg | HA |
| **Wnt7a_Δ1-149** | GGCSADIRYGIGFAKVFVDAREIKQNARTLMNLHNNEAGRKI LEENMKLECKCHGVSGSCTTKTCWTTLPQFRELGYVLKDKY NEAVHVE**PVRASRNKRPTFLKIKKP**LSYRKPMDTDLVYIEKS | aaggatccaccatgggtggctgctctgccgacatc | aagaattctcaagcgtaatctggaacatcgtatgggtacttgcacgtgtacatctccg | HA |
| **Wnt7a_Δ3aa*GSG** | MNRKARRCLGHLFLSLGMVYLRIGGFSSVVALGASIICNKIP GLAPRQRAICQSRPDAIIVIGEGSQMGLDECQFQFRNGRWN CSALGERTVFGKELKVGSREAAFTYAIIAAGVAHAITAACTQ GNLSDCGCDKEKQGQYHRDEGWKWGGCSADIRYGIGFAK VFVDAREGSGNARTLMNLHNNEAGRKILEENMKLECKCHGV | caaggtctttgtggatgcccgggagggctcggggaatgcccggactctcatgaacttgcac | gtgcaagttcatgagagtccgggcattccccgagccctcccgggcatccacaaagaccttg | HA |
| **Wnt7a_Δ3aa*ESP** | MNRKARRCLGHLFLSLGMVYLRIGGFSSVVALGASIICNKIP GLAPRQRAICQSRPDAIIVIGEGSQMGLDECQFQFRNGRWN CSALGERTVFGKELKVGSREAAFTYAIIAAGVAHAITAACTQ GNLSDCGCDKEKQGQYHRDEGWKWGGCSADIRYGIGFAK VFVDARE**PVRASRNKRPTFLKIKKP** | tggcttcttgatcttcaggaaggtgggccgcttgttgcggctggcacgcacaggctcccgggcatccacaaagaccttg | cctgtgcgtgccagccgcaacaagcggcccaccttcctgaagatcaagaagccaaatgcccggactctcatgaacttgc | HA |
| **Wnt7a_Δ213-349*ESP@172** | MNRKARRCLGHLFLSLGMVYLRIGGFSSVVALGASIICNKIP GLAPRQRAICQSRPDAIIVIGEGSQMGLDECQFQFRNGRWN CSALGERTVFGKELKVGSREAAFTYAIIAAGVAHAITAACTQ GNLSDCGCDKEKQGQYHRDEGWKWGGCSADIRYGIGFAK | aaggatccaccatgaaccggaaagcgcggcgctg | aagaattctcaagcgtaatctggaacatcgtatgggtacttggtggtgcacgagcctgac | HA |
| **Wnt7a_Δ213-349*ESP** | MNRKARRCLGHLFLSLGMVYLRIGGFSSVVALGASIICNKIP GLAPRQRAICQSRPDAIIVIGEGSQMGLDECQFQFRNGRWN CSALGERTVFGKELKVGSREAAFTYAIIAAGVAHAITAACTQ GNLSDCGCDKEKQGQYHRDEGWKWGGCSADIRYGIGFAK | aaggatccaccatgaaccggaaagcgcggcgctg | 1-gcttgttgcggctggcacgcacaggtcccgatcccttggtggtgcacgagcctgacac 2-agaattctcaagc | HA |
| **Wnt7a_*ESPΔ213-349** | **PVRASRNKRPTFLKIKKP**MNRKARRCLGHLFLSLGMVYLRIG GFSSVVALGASIICNKIPGLAPRQRAICQSRPDAIIVIGEGSQ MGLDECQFQFRNGRWNCSALGERTVFGKELKVGSREAAFT YAIIAAGVAHAITAACTQGNLSDCGCDKEKQGQYHRDEGWK | 1-cccaccttcctgaagatcaagaagccaggatcgggaatgaaccggaaagcgcggcgctg 2- | aagaattctcaagcgtaatctggaacatcgtatgggtacttggtggtgcacgagcctgac | HA |

| | | | | |
|---|---|---|---|---|
| **HALO*ESP-HA** | MAEIGTGFPFDPHYVEVLGERMHYVDVGPRDGTPVLFLHGN<br>PTSSYVWRNIIPHVAPTHRCIAPDLIGMGKSDKPDLGYFFDD<br>HVRFMDAFIEALGLEEVVLVIHDWGSALGFHWAKRNPERVK<br>GIAFMEFIRPIPTWDEWPEFARETFQAFRTTDVGRKLIIDQNV<br>FIEGTLPMGVVRPLTEVEMDHYREPFLNPVDREPLWRFPNE | atat aagctt acc atg atataagctt<br>atggaggatctgtactttcag | caagcggcccaccttcctgaagatc<br>aagaagccatac cca tac gat<br>gtt cca gat tac gct tga<br>gaattctt | HA |
| **HALO*ESP** | MAEIGTGFPFDPHYVEVLGERMHYVDVGPRDGTPVLFLHGN<br>PTSSYVWRNIIPHVAPTHRCIAPDLIGMGKSDKPDLGYFFDD<br>HVRFMDAFIEALGLEEVVLVIHDWGSALGFHWAKRNPERVK<br>GIAFMEFIRPIPTWDEWPEFARETFQAFRTTDVGRKLIIDQNV<br>FIEGTLPMGVVRPLTEVEMDHYREPFLNPVDREPLWRFPNE | atat aagctt acc atg atataagctt<br>atggaggatctgtactttcag | cttcaggaaggtgggccgcttgttgc<br>ggctggcacgcacaggtcccgatc<br>caccggaaatctccagagtag | N/A |
| **Wnt7a-BirA-myc** | MNRKARRCLGHLFFSLGMVYLRIGGFSSVVALGASIICNKIP<br>GLAPRQRAICQSRPDAIIVIGEGSQMGLDECQFQFRNGRWN<br>CSALGERTVFGKELKVGSREAAFTYAIIAAGVAHAITAACTQ<br>GNLSDCGCDKEKQGQYHRDEGWKWGGCSADIRYGIGFAK<br>VFVDAREIKQNARTLMNLHNNEAGRKILEENMKLECKCHGV<br>SGSCTTKTCWTTLPQFRELGYVLKDKYNEAVHVEPVRASRN<br>KRPTFLKIKKPLSYRKPMDTDLVYIEKSPNYCEEDPVTGSVG<br>TQGRACNKTAPQASGCDLMCCGRGYNTHQYARVWQCNCK | **1-**<br>tatagaattcgccaccatgaaccggaa<br>agcgcggcgctgcc<br>**2-**<br>tataaccggtggaagtggaagtggaagt<br>ggaagtgacttcaagaacctgatctggc<br>tg | **1-**<br>tatagtcgaccatatgtatataaccg<br>gtcttgcacgtgtacatctccgtgcg<br>c **2-**<br>tatacatatgtcaaagatcttcctcgg<br>atatgagtttctgctcctcgaggcttct<br>tctcaggctg | MYC |
| **Myc-BirA-ESP** | MEQKLISEEDLDFKNLIWLKEVDSTQERLKEWNVSYGTALV<br>ADRQTKGRGGLGRKWLSQEGGLYFSFLLNPKEFENLLQLPL<br>VLGLSVSEALEEITEIPFSLKWPNDVYFQEKKVSGVLCELSK<br>DKLIVGIGINVNQREIPEEIKDRATTLYEITGKDWDRKEVLLK<br>[illegible] | tatactcgagggatcgggacctgtgcgt<br>gccagccgcaac | gtcgactcatggcttcttgatcttcag<br>gaag | MYC |
| **Wnt7a_ESP*Scramb** | MNRKARRCLGHLFLSLGMVYLRIGGFSSVVALGASIICNKIP<br>GLAPRQRAICQSRPDAIIVIGEGSQMGLDECQFQFRNGRWN<br>CSALGERTVFGKELKVGSREAAFTYAIIAAGVAHAITAACTQ<br>GNLSDCGCDKEKQGQYHRDEGWKWGGCSADIRYGIGFAK<br>VFVDAREIKQNARTLMNLHNNEAGRKILEENMKLECKCHGV | tggcgagcccgcgcattacctttaaacc<br>gaaacgccgcgtgctgtcgtaccgcaa<br>gcccatg | ggtttaaaggtaatgcgcgggctcgc<br>cagtttttttgttcgGctccacgtgaac<br>ggcctcgttg | HA |
| **Wnt7a_K247A** | MNRKARRCLGHLFLSLGMVYLRIGGFSSVVALGASIICNKIP<br>GLAPRQRAICQSRPDAII<br>VIGEGSQMGLDECQFQFRNGRWNCSALGERTVFGKELKVG<br>SREAAFTYAIIAAGVAHAIT<br>AACTQGNLSDCGCDKEKQGQYHRDEGWKWGGCSADIRYG | gtgcgtgccagccgcaacgcgcggc<br>ccaccttcctgaagatc | gatcttcaggaaggtgggccgcgcg<br>ttgcggctggcacgcac | HA |
| **Wnt7a_K253A** | MNRKARRCLGHLFLSLGMVYLRIGGFSSVVALGASIICNKIP<br>GLAPRQRAICQSRPDAII<br>VIGEGSQMGLDECQFQFRNGRWNCSALGERTVFGKELKVG<br>SREAAFTYAIIAAGVAHAIT<br>AACTQGNLSDCGCDKEKQGQYHRDEGWKWGGCSADIRYG | caagcggcccaccttcctgGCgatca<br>agaagccactgtcgtac | gtacgacagtggcttcttgatcGC<br>caggaaggtgggccgcttg | HA |

| | | | | |
|---|---|---|---|---|
| **Wnt7a_K255A** | MNRKARRCLGHLFLSLGMVYLRIGGFSSVVALGASIICNKIP<br>GLAPRQRAICQSRPDAII<br>VIGEGSQMGLDECQFQFRNGRWNCSALGERTVFGKELKVG<br>SREAAFTYAIIAAGVAHAIT<br>AACTQGNLSDCGCDKEKQGQYHRDEGWKWGGCSADIRYG | cggcccaccttcctgaagatcGCgaa<br>gccactgtcgtaccgcaag | cttgcggtacgacagtggcttcgcg<br>atcttcaggaaggtgggccg | HA |
| **Wnt7a_K256A** | MNRKARRCLGHLFLSLGMVYLRIGGFSSVVALGASIICNKIP<br>GLAPRQRAICQSRPDAII<br>VIGEGSQMGLDECQFQFRNGRWNCSALGERTVFGKELKVG<br>SREAAFTYAIIAAGVAHAIT<br>AACTQGNLSDCGCDKEKQGQYHRDEGWKWGGCSADIRYG<br>IGFAKVFVDAREIKQNARTLM | caccttcctgaagatcaagGCgccact<br>gtcgtaccgcaagccc | gggcttgcggtacgacagtggcgc<br>cttgatcttcaggaaggtg | HA |
| **Wnt7a_ΔESP*Wnt10a-ESP** | MNRKARRCLGHLFLSLGMVYLRIGGFSSVVALGASIICNKIP<br>GLAPRQRAICQSRPDAII<br>VIGEGSQMGLDECQFQFRNGRWNCSALGERTVFGKELKVG<br>SREAAFTYAIIAAGVAHAIT<br>AACTQGNLSDCGCDKEKQGQYHRDEGWKWGGCSADIRYG | 1-<br>aaggatccaccatgaaccggaaagcg<br>cggcgctg<br>2-gct ccg ggc gct ccc ggg ccg<br>cgc cga cgg gcc agc | 1-<br>tgggcccggctccagctggccgcc<br>gttgcggttgtgaggctccacgtgaa<br>cggcctc<br>2- | HA |
| **Wnt7a_ΔESP*Wnt16-ESP** | MNRKARRCLGHLFLSLGMVYLRIGGFSSVVALGASIICNKIP<br>GLAPRQRAICQSRPDAII<br>VIGEGSQMGLDECQFQFRNGRWNCSALGERTVFGKELKVG<br>SREAAFTYAIIAAGVAHAIT<br>AACTQGNLSDCGCDKEKQGQYHRDEGWKWGGCSADIRYG | 1-<br>aaggatccaccatgaaccggaaagcg<br>cggcgctg2-<br>agg aga gaa aaa gat cag agg<br>aaa ata cca atc cat gacacgga | 1-<br>atctttttctctcctgcgcattttcctctt<br>tgttttaggctccacgtgaacggcct<br>c 2-<br>aagaattctcaagcgtaatctggaac | HA |
| **Wnt10b_ΔESP*GSGS** | MLEEPRPRPPPSGLAGLLFLALCSRALSNEILGLKLPGEPPL<br>TANTVCLTLSGLSKRQLG<br>LCLRNPDVTASALQGLHIAVHECQHQLRDQRWNCSALEGG<br>GRLPHHSAILKRGFRESAFS<br>FSMLAAGVMHAVATACSLGKLVSCGCGWKGSGEQDRLRA<br>KLLQLQALSRGKSFPHSLPSP | 1-<br>aaggatccaccatgctggaggagccc<br>cggcc<br>2-cgg ctg ggc cgg gcc atc ttc<br>att ggttcaggttcagag ctg gtc<br>tac ttt gag aag tct c | 1- g aga ctt ctc aaa gta gac<br>cag ctctgaacctgaacc aat<br>gaa gat ggc ccg gcc cag<br>ccg 2-<br>aagaattcctaaccggtacgcgtag<br>aatcg | HA |
| **Wnt10b_RR302AA** | MLEEPRPRPPPSGLAGLLFLALCSRALSNEILGLKLPGEPPL<br>TANTVCLTLSGLSKRQLG<br>LCLRNPDVTASALQGLHIAVHECQHQLRDQRWNCSALEGG<br>GRLPHHSAILKRGFRESAFS<br>FSMLAAGVMHAVATACSLGKLVSCGCGWKGSGEQDRLRA<br>KLLQLQALSRGKSFPHSLPSP<br>GPGSSPSPGPQDTWEWGGCNHDMDFGEKFSRDFLDSREA<br>PRDIQARMRIHNNRVGRQVVT | gcc ttc cag ccc cgt ctg cgt<br>ccc gct gcc ctc tca gga gag ct<br>g gtc tac ttt g | c aaa gta gac cag ctc tcc<br>tga gag ggc agc gggacg cag<br>acg ggg ctg gaa ggc | HA |
