## Supplemental Figures 1-9 for "Wnt binding to Coatomer proteins directs secretion on exosomes independently of palmitoylation"

#### Extended Data Fig.1

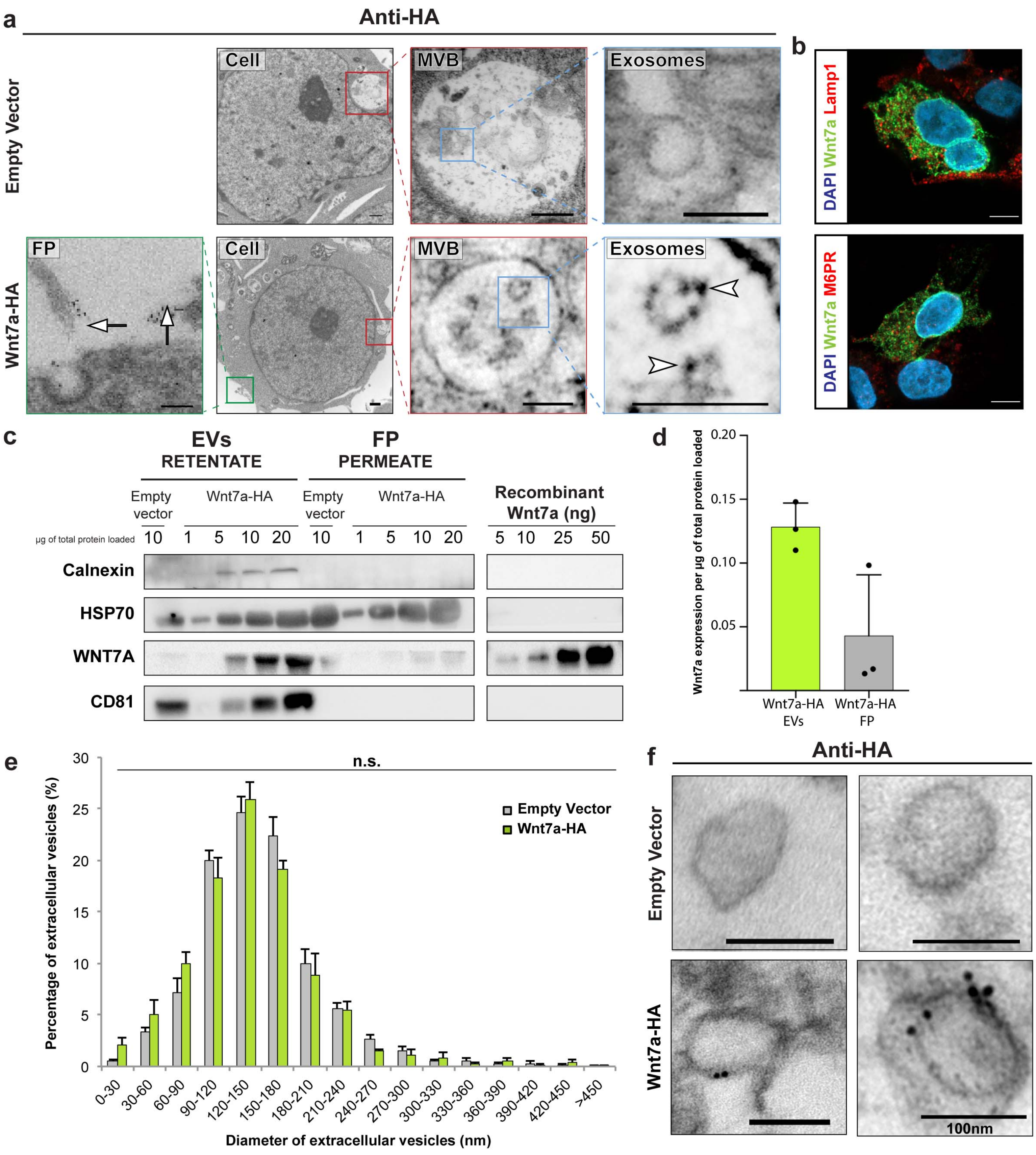

Extended Data Fig.2

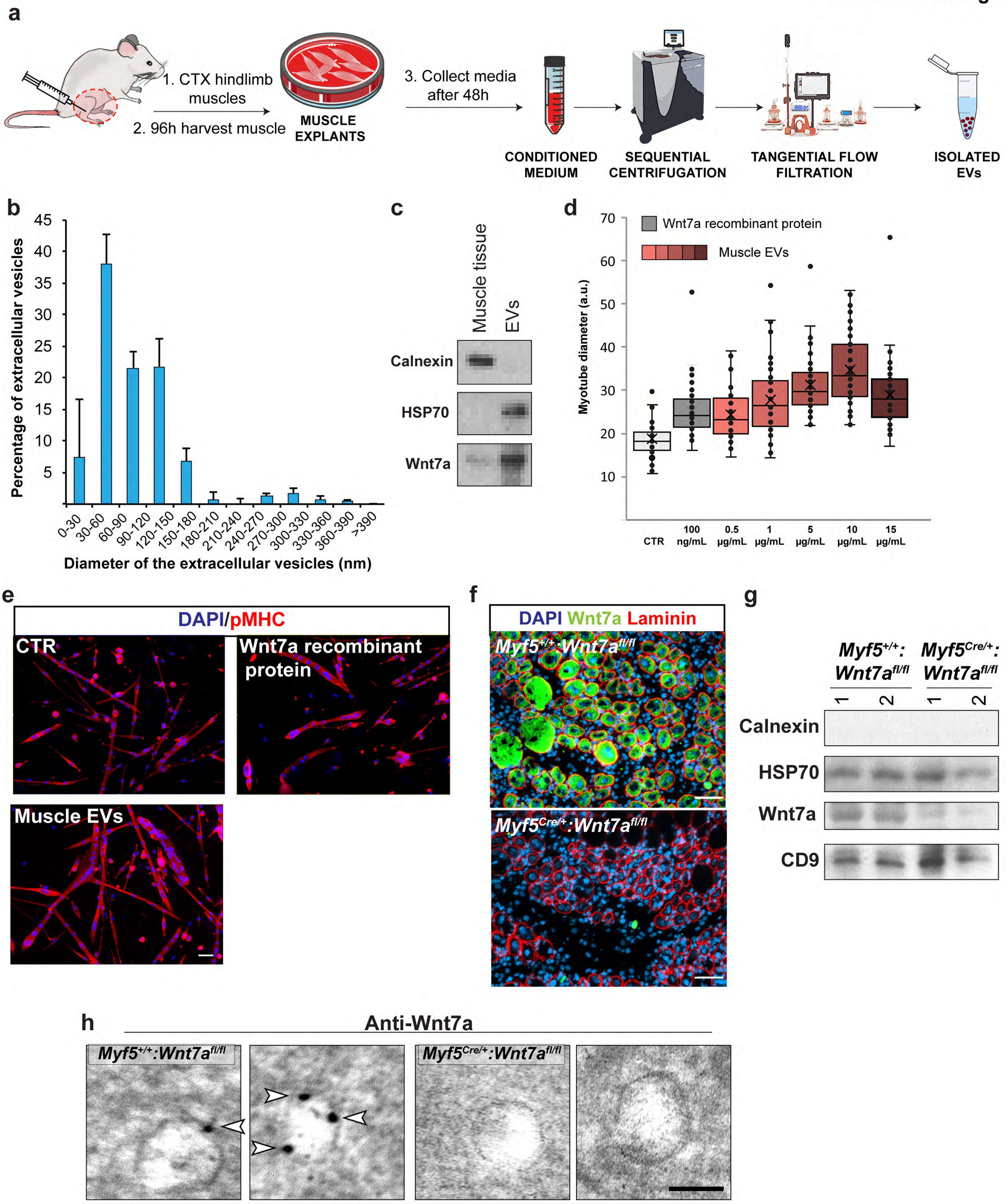

Extended Data Fig.3

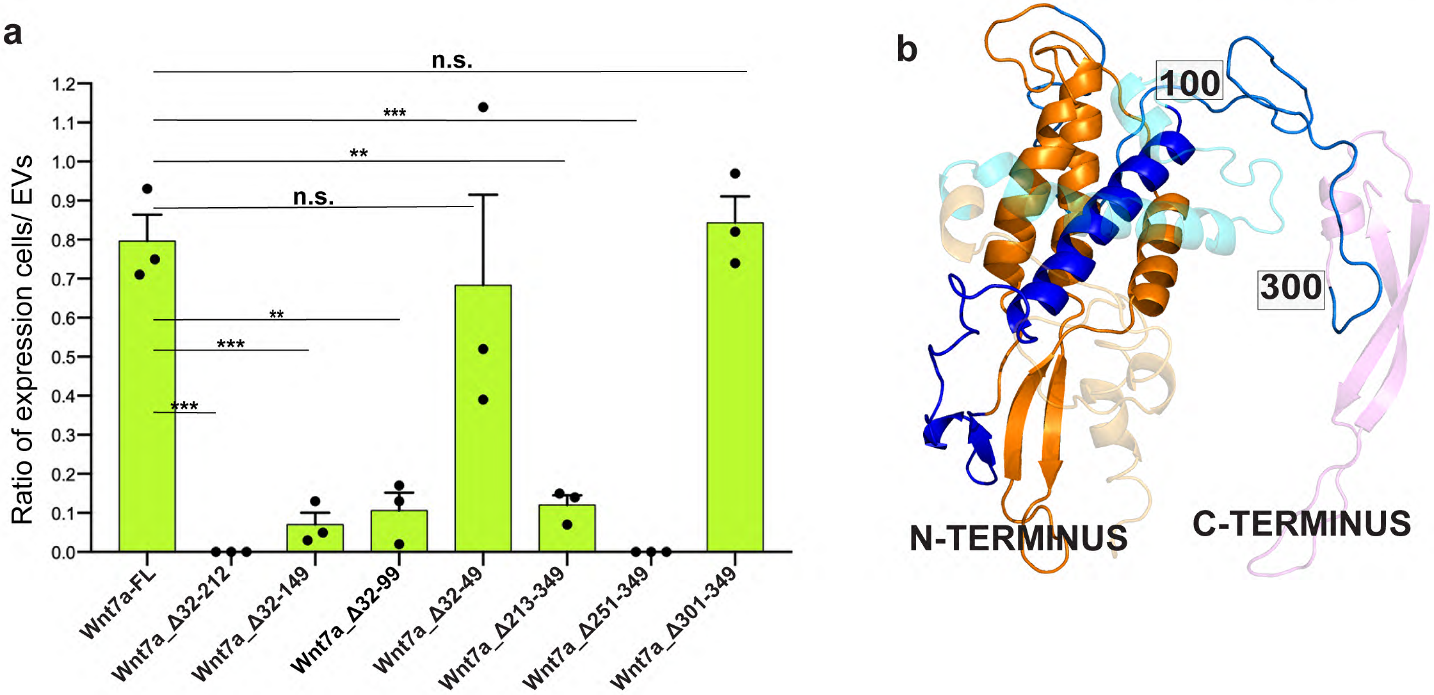

### Extended Data Fig.4

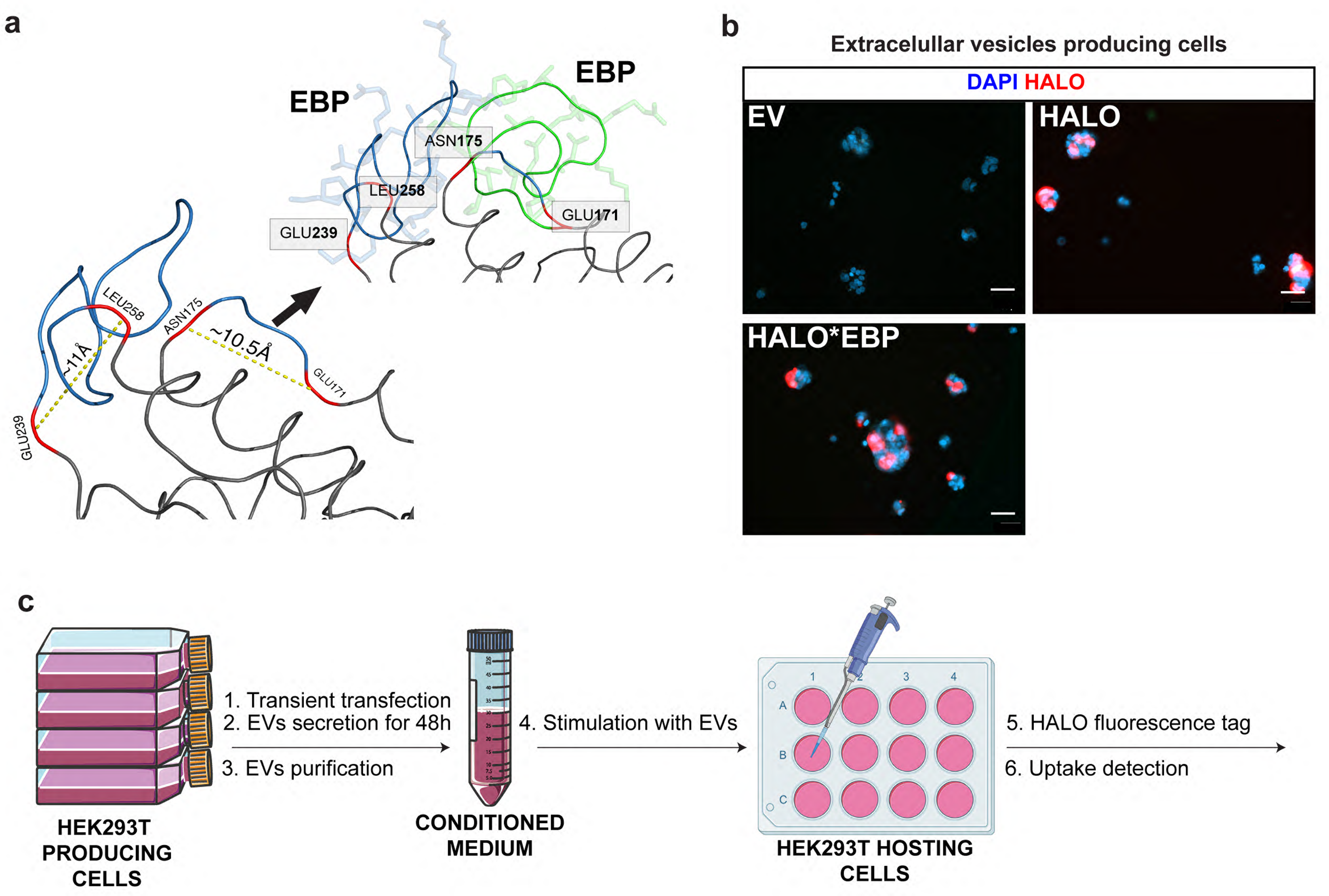

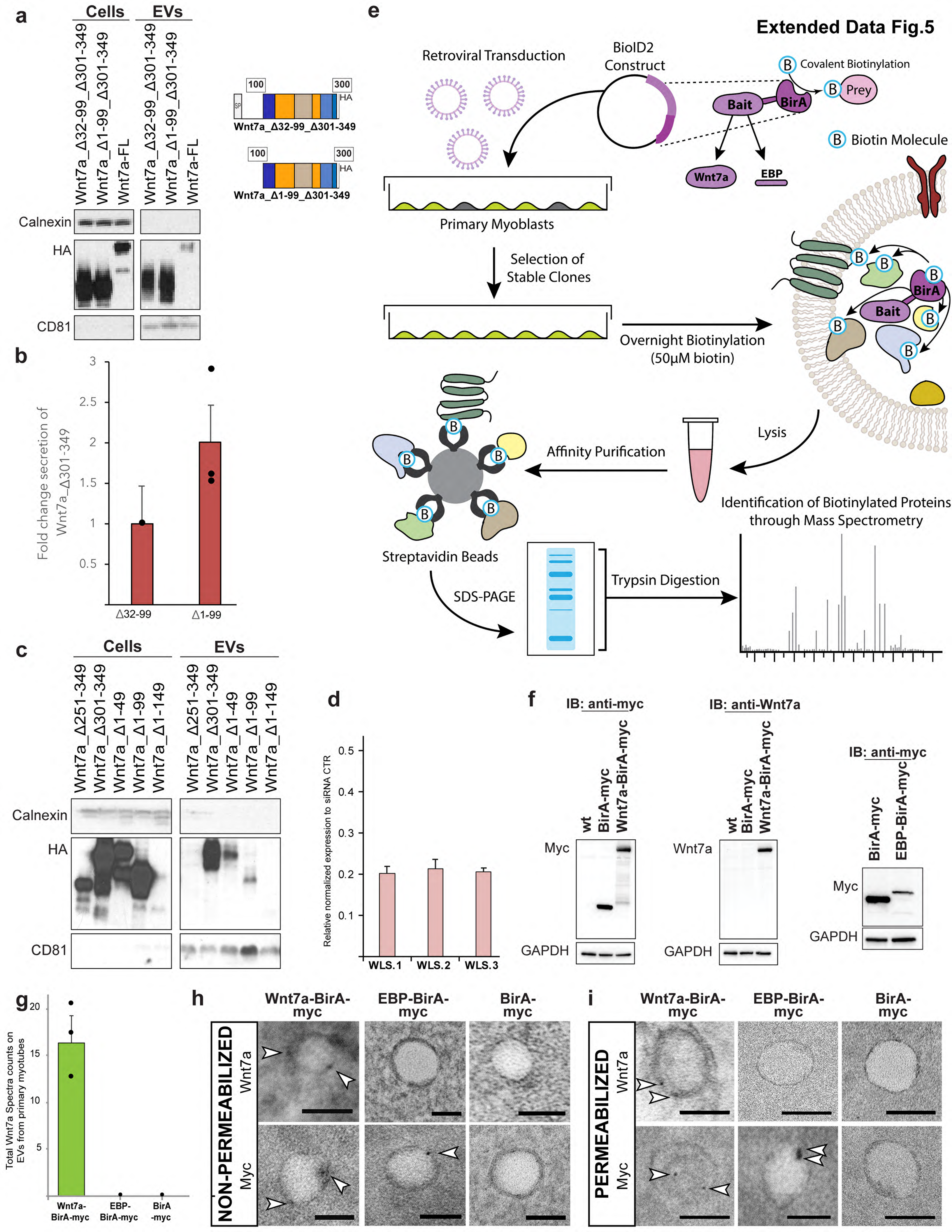

Extended Data Fig.5
a
Cells
EVs
Wnt7a_Δ32-99_Δ301-349
Wnt7a_Δ1-99_Δ301-349
Wnt7a-FL
Calnexin
HA
CD81
100
300
SP
HA
Wnt7a_Δ32-99_Δ301-349
Wnt7a_Δ1-99_Δ301-349
b
Fold change secretion of Wnt7a_Δ301-349
Δ32-99
Δ1-99
c
Wnt7a_Δ251-349
Wnt7a_Δ301-349
Wnt7a_Δ1-49
Wnt7a_Δ1-99
Wnt7a_Δ1-149
d
Relative normalized expression to siRNA CTR
WLS.1
WLS.2
WLS.3
e
Retroviral Transduction
BioID2 Construct
Covalent Biotinylation
Bait
BirA
Prey
Biotin Molecule
Wnt7a
EBP
Primary Myoblasts
Selection of Stable Clones
Overnight Biotinylation (50μM biotin)
Lysis
Affinity Purification
Streptavidin Beads
SDS-PAGE
Trypsin Digestion
Identification of Biotinylated Proteins through Mass Spectrometry
f
IB: anti-myc
IB: anti-Wnt7a
wt
BirA-myc
Wnt7a-BirA-myc
EBP-BirA-myc
Myc
Wnt7a
GAPDH
g
Total Wnt7a Spectra counts on EVs from primary myotubes
Wnt7a-BirA-myc
EBP-BirA-myc
BirA-myc
h
i
NON-PERMEABILIZED
PERMEABILIZED
Wnt7a
Myc

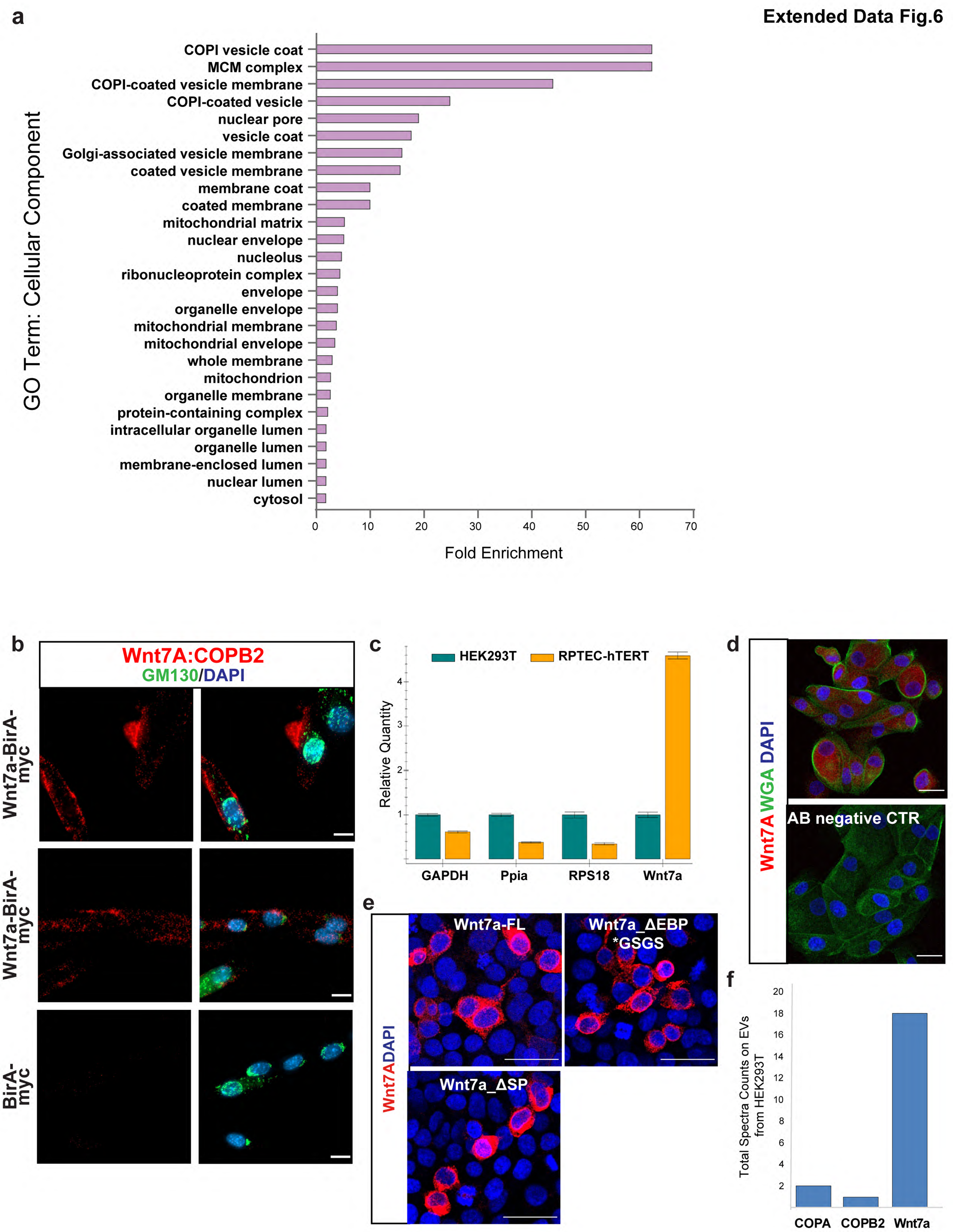
a
GO Term: Cellular Component
COPI vesicle coat
MCM complex
COPI-coated vesicle membrane
COPI-coated vesicle
nuclear pore
vesicle coat
Golgi-associated vesicle membrane
coated vesicle membrane
membrane coat
coated membrane
mitochondrial matrix
nuclear envelope
nucleolus
ribonucleoprotein complex
envelope
organelle envelope
mitochondrial membrane
mitochondrial envelope
whole membrane
mitochondrion
organelle membrane
protein-containing complex
intracellular organelle lumen
organelle lumen
membrane-enclosed lumen
nuclear lumen
cytosol
0 10 20 30 40 50 60 70
Fold Enrichment
b
Wnt7A:COPB2
GM130/DAPI
Wnt7a-BirA-myc
Wnt7a-BirA-myc
BirA-myc
c
HEK293T
RPTEC-hTERT
Relative Quantity
1 2 3 4
GAPDH
Ppia
RPS18
Wnt7a
d
Wnt7A WGA DAPI
AB negative CTR
e
Wnt7ADAPI
Wnt7a-FL
Wnt7a_ΔEBP *GSGS
Wnt7a_ΔSP
f
Total Spectra Counts on EVs from HEK293T
2 4 6 8 10 12 14 16 18 20
COPA
COPB2
Wnt7a

**Extended Data Fig.7**

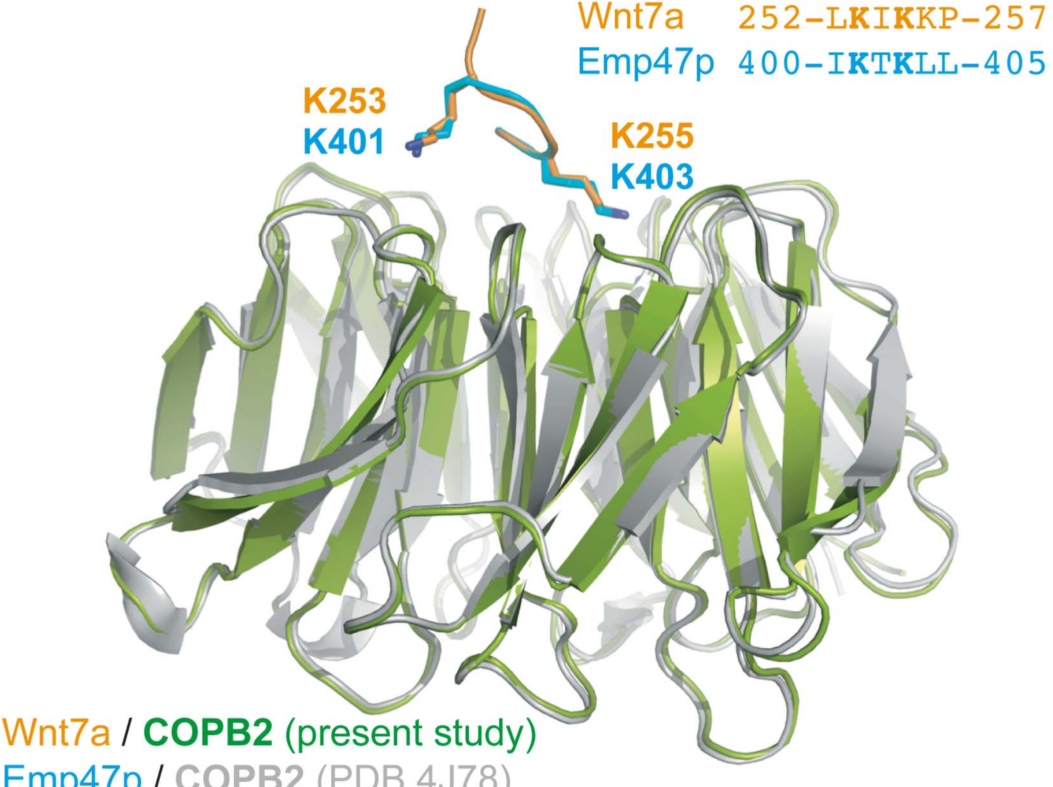

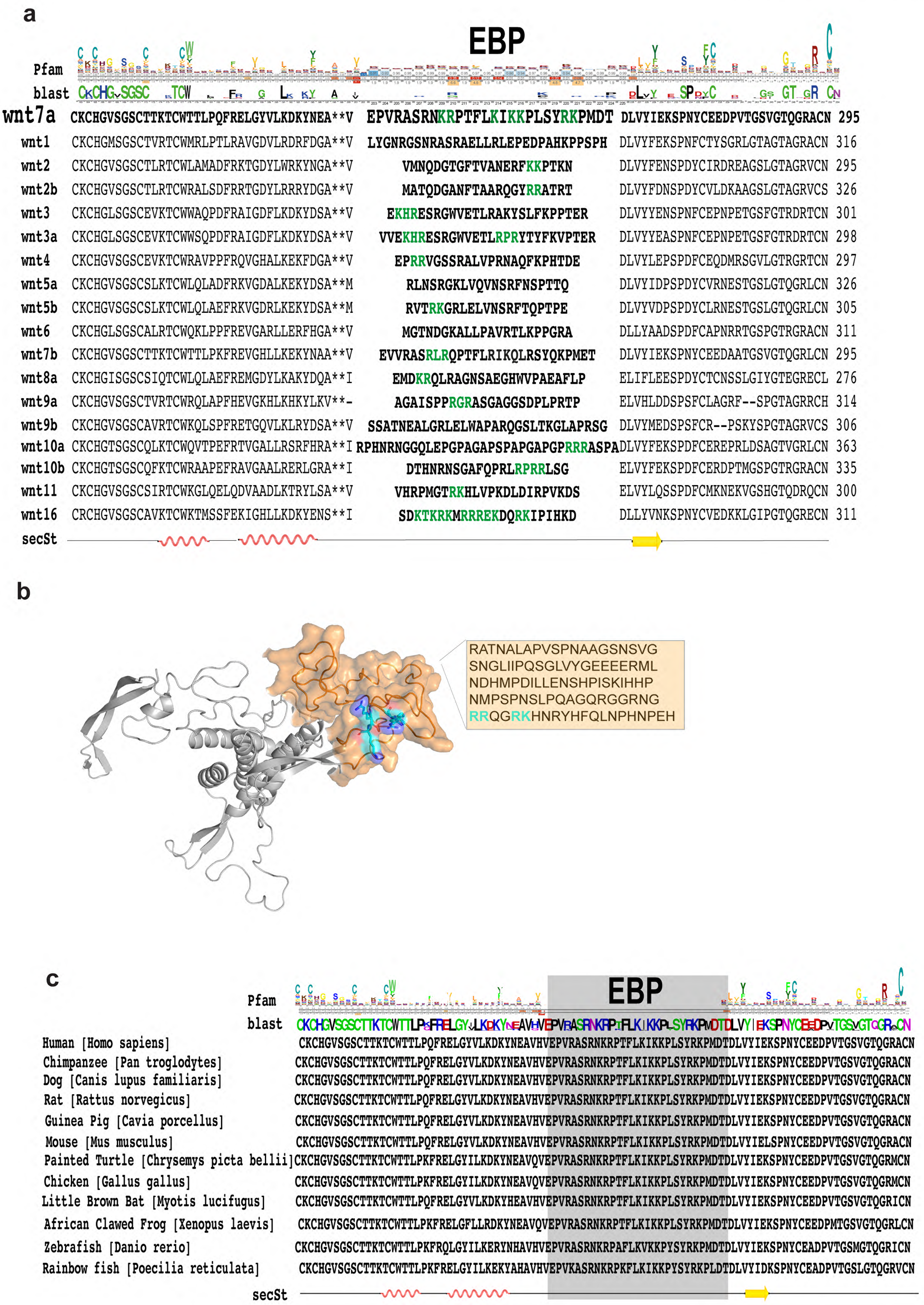
a
EBP
Pfam
blast
wnt7a CKCHGVSGSCTTKTCWTTLPQFRELGYVLKDKYNEA**V EPVRASRNKRPTFLKIKKPLSYRKPMDT DLVYIEKSPNYCEEDPVTGSVGTQGRACN 295
wnt1 CKCHGMSGSCTVRTCWMRLPTLRAVGDVLRDRFDGA**V LYGNRGSNRASRAELLRLEPEDPAHKPPSPH DLVYFEKSPNFCTYSGRLGTAGTAGRACN 316
wnt2 CKCHGVSGSCTLRTCWLAMADFRKTGDYLWRKYNGA**V VMNQDGTGFTVANERFKKPTKN DLVYFENSPDYCIRDREAGSLGTAGRVCN 295
wnt2b CKCHGVSGSCTLRTCWRALSDFRRTGDYLRRRYDGA**V MATQDGANFTAARQGYRRATRT DLVYFDNSPDYCVLDKAAGSLGTAGRVCS 326
wnt3 CKCHGLSGSCEVKTCWWAQPDFRAIGDFLKDKYDSA**V EKHRESRGWVETLRAKYSLFKPPTER DLVYYENSPNFCEPNPETGSFGTRDRTCN 301
wnt3a CKCHGLSGSCEVKTCWWSQPDFRAIGDFLKDKYDSA**V VVEKHRESRGWVETLRPRYTYFKVPTER DLVYYEASPNFCEPNPETGSFGTRDRTCN 298
wnt4 CKCHGVSGSCEVKTCWRAVPPFRQVGHALKEKFDGA**V EPRRVGSSRALVPRNAQFKPHTDE DLVYLEPSPDFCEQDMRSGVLGTRGRTCN 297
wnt5a CKCHGVSGSCSLKTCWLQLADFRKVGDALKEKYDSA**M RLNSRGKLVQVNSRFNSPTTQ DLVYIDPSPDYCVRNESTGSLGTQGRLCN 326
wnt5b CKCHGVSGSCSLKTCWLQLAEFRKVGDRLKEKYDSA**M RVTRKGRLELVNSRFTQPTPE DLVYVDPSPDYCLRNESTGSLGTQGRLCN 305
wnt6 CKCHGLSGSCALRTCWQKLPPFREVGARLLERFHGA**V MGTNDGKALLPAVRTLKPPGRA DLLYAADSPDFCAPNRRTGSPGTRGRACN 311
wnt7b CKCHGVSGSCTTKTCWTTLPKFREVGHLLKEKYNAA**V EVVRASRLRQPTFLRIKQLRSYQKPMET DLVYIEKSPNYCEEDAATGSVGTQGRLCN 295
wnt8a CKCHGISGSCSIQTCWLQLAEFREMGDYLKAKYDQA**I EMDKRQLRAGNSAEGHWVPAEAFLP ELIFLEESPDYCTCNSSLGIYGTEGRECL 276
wnt9a CKCHGVSGSCTVRTCWRQLAPFHEVGKHLKHKYLKV**- AGAISPPRGRASGAGGSDPLPRTP ELVHLDDSPSFCLAGRF--SPGTAGRRCH 314
wnt9b CKCHGVSGSCAVRTCWKQLSPFRETGQVLKLRYDSA**V SSATNEALGRLELWAPARQGSLTKGLAPRSG DLVYMEDSPSFCR--PSKYSPGTAGRVCS 306
wnt10a CKCHGTSGSCQLKTCWQVTPEFRTVGALLRSRFHRA**I RPHNRNGGQLEPGPAGAPSPAPGAPGPRRRASPA DLVYFEKSPDFCEREPRLDSAGTVGRLCN 363
wnt10b CKCHGTSGSCQFKTCWRAAPEFRAVGAALRERLGRA**I DTHNRNSGAFQPRLRPRRLSG ELVYFEKSPDFCERDPTMGSPGTRGRACN 335
wnt11 CKCHGVSGSCSIRTCWKGLQELQDVAADLKTRYLSA**V VHRPMGTRKHLVPKDLDIRPVKDS ELVYLQSSPDFCMKNEKVGSHGTQDRQCN 300
wnt16 CRCHGVSGSCAVKTCWKTMSSFEKIGHLLKDKYENS**I SDKTKRKMRRREKDQRKIPIHKD DLLYVNKSPNYCVEDKKLGIPGTQGRECN 311
secSt
b
RATNALAPVSPNAAGSNSVG
SNGLIIPQSGLVYGEEEERML
NDHMPDILLENSHPISKIHHP
NMPSPNSLPQAGQRGGRNG
RRQGRKHNRYHFQLNPHNPEH
c
EBP
Pfam
blast
Human [Homo sapiens] CKCHGVSGSCTTKTCWTTLPQFRELGYVLKDKYNEAVHVEPVRASRNKRPTFLKIKKPLSYRKPMDTDLVYIEKSPNYCEEDPVTGSVGTQGRACN
Chimpanzee [Pan troglodytes] CKCHGVSGSCTTKTCWTTLPQFRELGYVLKDKYNEAVHVEPVRASRNKRPTFLKIKKPLSYRKPMDTDLVYIEKSPNYCEEDPVTGSVGTQGRACN
Dog [Canis lupus familiaris] CKCHGVSGSCTTKTCWTTLPQFRELGYVLKDKYNEAVHVEPVRASRNKRPTFLKIKKPLSYRKPMDTDLVYIEKSPNYCEEDPVTGSVGTQGRACN
Rat [Rattus norvegicus] CKCHGVSGSCTTKTCWTTLPQFRELGYVLKDKYNEAVHVEPVRASRNKRPTFLKIKKPLSYRKPMDTDLVYIEKSPNYCEEDPVTGSVGTQGRACN
Guinea Pig [Cavia porcellus] CKCHGVSGSCTTKTCWTTLPQFRELGYVLKDKYNEAVHVEPVRASRNKRPTFLKIKKPLSYRKPMDTDLVYIEKSPNYCEEDPVTGSVGTQGRACN
Mouse [Mus musculus] CKCHGVSGSCTTKTCWTTLPQFRELGYVLKDKYNEAVHVEPVRASRNKRPTFLKIKKPLSYRKPMDTDLVYIELSPNYCEEDPVTGSVGTQGRACN
Painted Turtle [Chrysemys picta bellii] CKCHGVSGSCTTKTCWTTLPKFRELGYILKDKYNEAVQVEPVRASRNKRPTFLKIKKPLSYRKPMDTDLVYIEKSPNYCEEDPVTGSVGTQGRMCN
Chicken [Gallus gallus] CKCHGVSGSCTTKTCWTTLPKFRELGYILKDKYNEAVQVEPVRASRNKRPTFLKIKKPLSYRKPMDTDLVYIEKSPNYCEEDPVTGSVGTQGRMCN
Little Brown Bat [Myotis lucifugus] CKCHGVSGSCTTKTCWTTLPQFRELGYVLKDKYHEAVHVEPVRASRNKRPTFLKIKKPLSYRKPMDTDLVYIEKSPNYCEEDPVTGSVGTQGRICN
African Clawed Frog [Xenopus laevis] CKCHGVSGSCTTKTCWTTLPKFRELGFLLRDKYNEAVQVEPVRASRNKRPTFLKIKKPLSYRKPMDTDLVYIEKSPNYCEEDPMTGSVGTQGRLCN
Zebrafish [Danio rerio] CKCHGVSGSCTTKTCWTTLPKFRQLGYILKERYNHAVHVEPVRASRNKRPAFLKVKKPYSYRKPMDTDLVYIEKSPNYCEADPVTGSMGTQGRICN
Rainbow fish [Poecilia reticulata] CKCHGVSGSCTTKTCWTTLPKFRELGYILKEKYAHAVHVEPVKASRNKRPKFLKIKKPYSYRKPLDTDLVYIDKSPNYCEADPVTGSLGTQGRVCN
secSt

Extended Data Fig.9

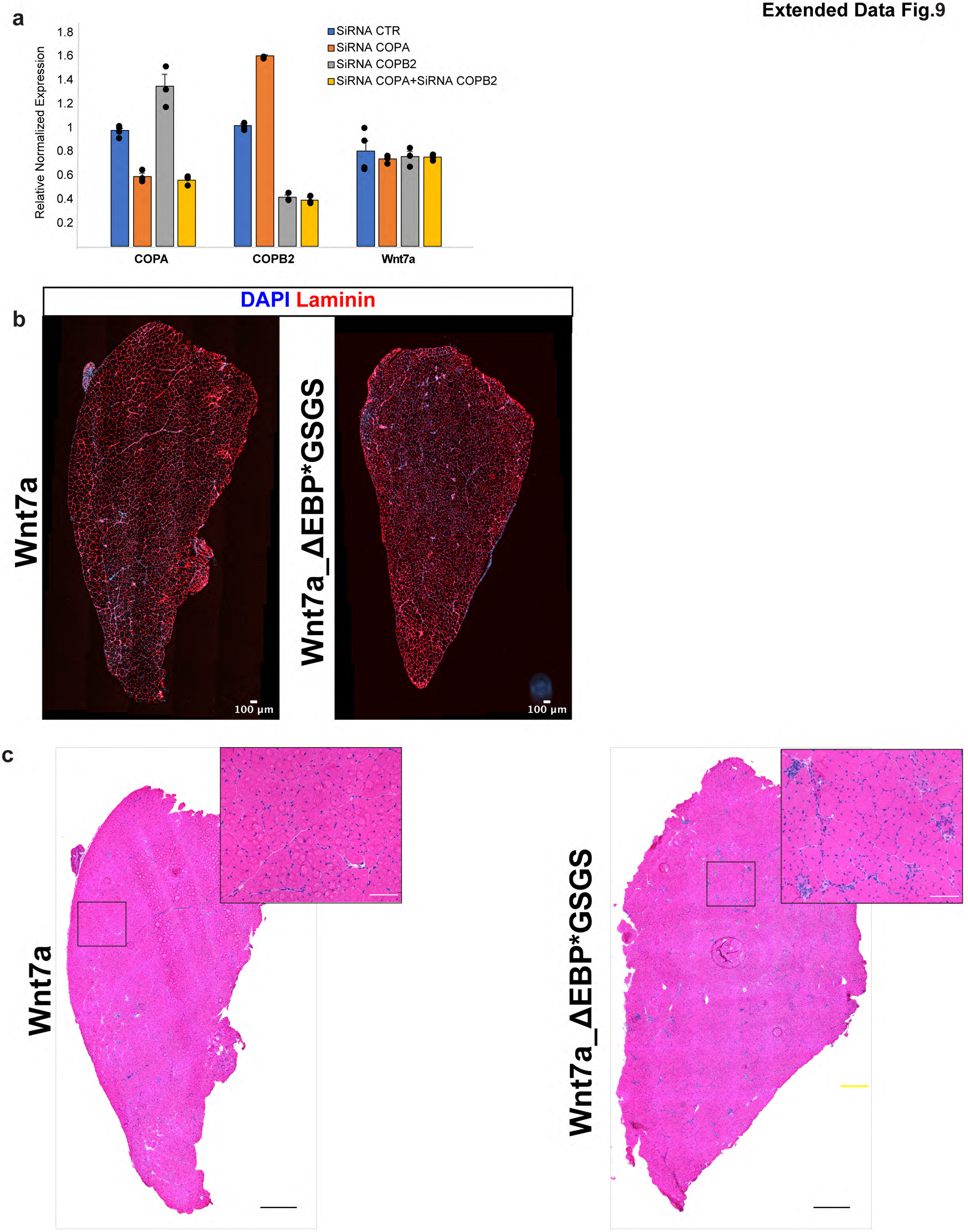
